# NANUQ^+^: A divide-and-conquer approach to network estimation

**DOI:** 10.1101/2024.10.30.621146

**Authors:** Elizabeth S. Allman, Hector Baños, John A. Rhodes, Kristina Wicke

## Abstract

Inference of a species network from genomic data remains a difficult problem, with recent progress mostly limited to the level-1 case. However, inference of the *Tree of Blobs* of a network, showing only the network’s cut edges, can be performed for any network by TINNiK, suggesting a divide-and-conquer approach to network inference where the tree’s multifurcations are individually resolved to give more detailed structure. Here we develop a method, NANUQ^+^, to quickly perform such a level-1 resolution. Viewed as part of the NANUQ pipeline for fast level-1 inference, this gives tools for both understanding when the level-1 assumption is likely to be met and for exploring all highly-supported resolutions to cycles.

## 1 Introduction

Inference of a phylogenetic species network from biological data is a challenging problem, with many theoretical and practical questions still unresolved. Formulated under the network multispecies coalescent model, such inference addresses two confounded sources of gene tree incongruence — reticulations in the network and incomplete lineage sorting.

Data for this is generally multilocus sequences of a genomic scale, with many approaches explicitly inferring gene trees for each locus as well as the species network (see [16] for an exception). Bayesian analyses are limited to very small data sets by the computational cost of simultaneous inference [27, 31]. Two-stage methods, that first infer individual gene trees and then treat them as data for network inference, scale better. However, even using a simplified pseudolikelihood as an optimality criterion and a heuristic search over network space, restricting the number of allowed reticulations and/or the level of the network is often necessary [24, 30].

The NANUQ and TINNiK algorithms [1, 2], on which this work builds, follow a two-stage approach but avoid a search over network space by instead computing certain intertaxon distances from quartet summaries of the gene trees. Assuming the network is level-1, the splits-graph of the expected NANUQ distance has a form that closely reflects the network structure. TINNiK makes no assumption on the network level, inferring only a *Tree of Blobs* showing just tree-like features of the network.

The empirical NANUQ distance of course has noise, and though this may be partially mitigated by the algorithm, for some data sets the splits-graph structure may not unambiguously support a single level-1 structure for the network. One goal of this work is to provide a quantitative method of choosing a specific level-1 structure in such ambiguous cases, thus allowing for automation of the last, interpretive step of the NANUQ algorithm [2]. NANUQ’s ability to visually suggest this ambiguity remains important, since it may expose a violation of the level-1 assumption. However, by quantifying support for various level-1 alternatives one may also address cases in which the ambiguity is suspected to arise mostly from noise. This work thus provides a theoretical analysis allowing for a measure of the fit of the data to level-1 structures, as well as providing a software implementation in the `MSCquartets` R package.

A second goal is to lay out an instance of a divide-and-conquer approach to network inference that we believe may be useful more broadly. We first infer the tree of blobs of the network, in which all reticulation cycles are collapsed to nodes. This tree might be obtained from the NANUQ splits-graph (assuming the network is level-1) or the TINNiK algorithm (for general networks). If we suspect a multifurcation arises from a cycle, we can focus on it individually and infer an optimal cycle structure. We do this with a least-squares approach, comparing an empirical NANUQ distance relating groups of taxa around the multifurcation to an expected one for each possible cycle resolution. This search can be done quickly for cycles of size less than about 10, and we offer a heuristic method for larger cycles, using a variant quartet-based distance. With multifurcations resolved individually, we can then combine these with the tree of blobs to obtain a more complete network structure, provided the resolutions collectively allow for a network rooting to exist. If a blob does not appear to be from a cycle, it may be left unresolved, so a global level-1 assumption is not needed if only some blobs appear to arise from cycles.

At this time we know of no other divide-and-conquer means of network inference, but we believe this approach is a promising one for further development. New methods for inferring the tree of blobs might be found, as well as new approaches for resolving its individual multifurcations under various assumptions on structure. For some types of blobs beyond cycles we suspect that future work will lead to consistent methods of inference of at least partial blob structure (see, e.g., [22]), which can then become part of an enhanced analysis pipeline. Other blobs may be sufficiently complex that data is insufficient to support a resolution.

This paper proceeds as follows: In the “Background“ section we lay out definitions and terminology. The next section, “Quartet distances“, develops a parametric family of distances that includes the one used in NANUQ, as well as a new variant useful in a fast heuristic for identifying large cycle structures. An algorithm for resolving individual multifurcations in a tree of blobs into cycles is developed in “Cycle resolution“. A section on “Simulated and empirical data analysis“ investigates the performance of these methods, as implemented in the R package `MSCquartets 3.0`, in simulations and with gene trees from a recent study of *Leopardus* [17], illustrating a full analysis process. This empirical analysis with additional commentary is also provided in a vignette in `MSCquartets`.

Note: As this project was being completed, we learned of independent work by Holtgrefe et al. [13] which uses quartet information to first obtain many candidate trees of blobs and then their level-1 network resolutions before choosing an optimal one. In that work a sum similar to our quartet distances for a blob is used, although with a different optimality criterion.

## 2 Background

Here we set notation and informally give definitions related to the multispecies coalescent model (NMSC) on a phylogenetic network, following those given more precisely in Allman et al. [2, 4]. Throughout, *X* = {*x*_1_, …, *x*_*n*_} denotes a non-empty finite set of taxa.

### 2.1 Rooted and unrooted phylogenetic networks

A *topological rooted phylogenetic network N* ^+^ is a finite, connected, directed graph with a degree-2 *root* node away from which all edges are directed, whose leaves are labelled by *taxa* in a set *X*. We focus on *binary* networks, where all internal nodes other than the root are either *hybrid* (with indegree 2 and outdegree 1) or *tree* (with indegree 1 and outdegree 2). Edges with a tree node or leaf child are tree edges, and edges with a hybrid child are hybrid edges.

To parameterize the NMSC model, *N* ^+^ must be a *metric network*. For this all edges have lengths (in coalescent units) and hybrid edges have *hybridization* or *inheritance probabilities*.

An *unrooted phylogenetic network* is obtained from a rooted one in several steps. First all edges and nodes above the *least stable ancestor*, LSA(*X*), a network generalization of the MRCA, are deleted. Then all tree edges are undirected, while hybrid edges retain their directions. Finally, LSA(*X*) is suppressed, with its incident edges conjoined (since we assume the network is binary). The resulting network *N* ^−^ is also referred to as a *semidirected phylogenetic network* in some works. A metric structure on *N* ^+^ is inherited by *N* ^−^ in the natural way.

Fig. 1 depicts a topological rooted binary phylogenetic network *N* ^+^ on the left and its induced topological unrooted phylogenetic network on the right. Finally, we use *N* to refer to a phylogenetic network that could either be unrooted or rooted.

**Fig. 1:**
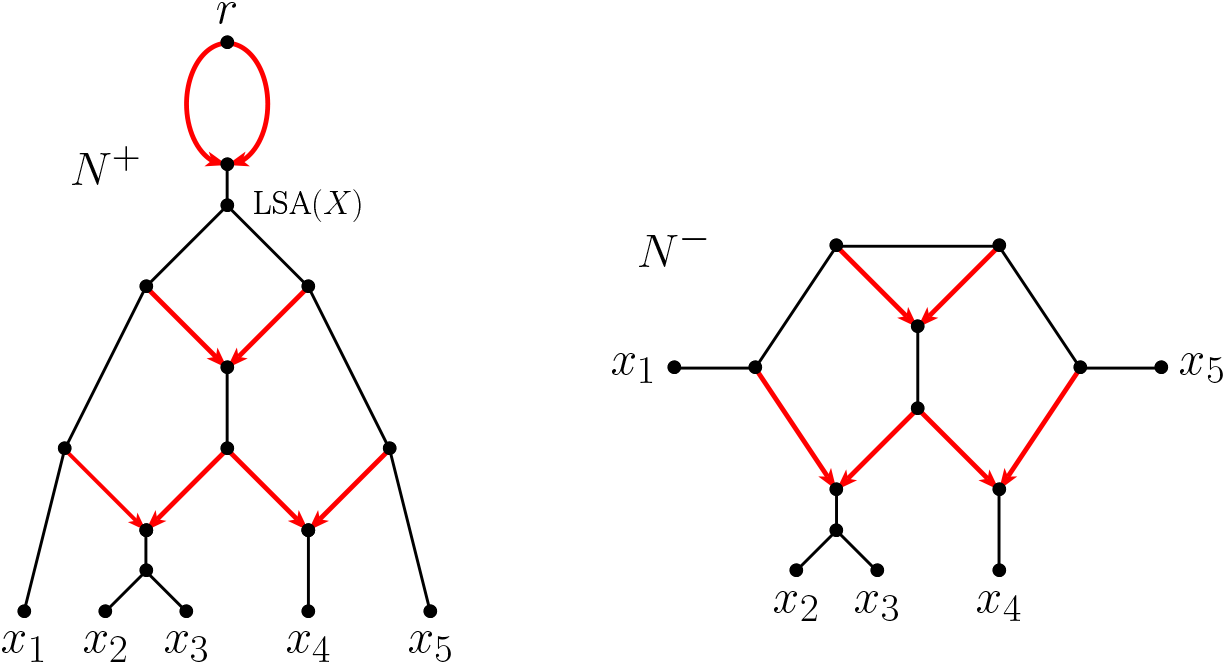
A topological rooted binary phylogenetic network *N* ^+^ (left) and its induced topological unrooted phylogenetic network *N* ^−^ (right). In *N* ^+^ only hybrid edges are highlighted as directed, but all other edges are implicitly assumed to be directed away from the root. In *N* ^−^ all non-hybrid edges are undirected.

Subnetworks of an unrooted network induced by sets of four taxa will play a central role in this work. For a precise definition, recall that a *trek* between vertices *x, y* in a network is the union of a semidirected path from some vertex *v* to *x* and a semidirected path from *v* to *y*. Such a trek is called *simple* (or an up-down path) if the two paths intersect only at *v*.

#### Definition 2.1.

Let *N* ^−^ be an unrooted phylogenetic network on *X* and let *x, y, z, w* ∈ *X*. The *induced quartet network Q*_*xyzw*_ is the unrooted network obtained by (1) keeping only the nodes and edges in simple treks between pairs of elements of {*x, y, z, w*}, and then (2) suppressing all degree-2 nodes.

That this definition yields the unrooted network obtained from the rooted quartet network induced from *N* ^+^ was shown in [7]. If *N* ^−^ is a metric network, the quartet network *Q*_*xyzw*_ naturally inherits a metric structure. Fig. 2 shows all quartet networks induced from the network *N* ^−^ of Fig. 1.

**Fig. 2:**
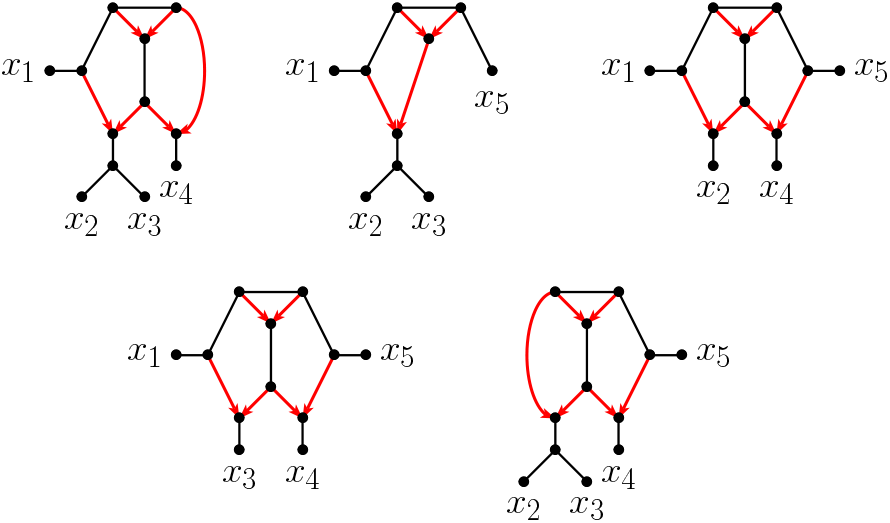
The five quartet networks induced from the topological unrooted phylogenetic network *N* ^−^ of Fig. 1.

We focus on the subclass of *level-1* phylogenetic networks. Since phylogenetic networks have no directed or semidirected cycles, we use the word *cycle* to refer to a sequence of edges which forms a cycle when all edges are undirected. For binary networks in particular, level-1 means no two cycles share a vertex, or equivalently, share an edge [23].

Within the class of level-1 networks, we first consider networks consisting of precisely one cycle with only leaves adjacent to it (i.e., there are no further tree-like structures).

#### Definition 2.2.

[11] An unrooted binary level-1 network *N* ^−^ on *X* with |*X*| = *n* is called an *n-sunlet* if it contains precisely one cycle of size *n*.

We sometimes call an *n*-sunlet an *n*-cycle network. The taxon adjacent to the (unique) hybrid node of a sunlet is referred to as the *hybrid taxon*. Additionally, we list taxa *x*_1_, …, *x*_*n*_, with *x*_1_ hybrid, proceeding consecutively around the cycle. We say taxa are *adjacent* to one another if they are consecutive in this order, also viewing *x*_*n*_ as adjacent to *x*_1_ (see Fig. 3).

**Fig. 3:**
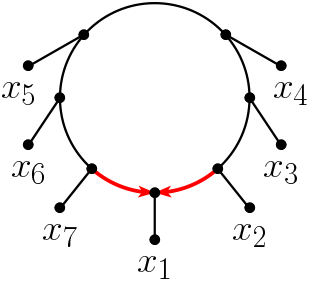
An unrooted 7-sunlet on *X* = {*x*_1_, …, *x*_7_} with hybrid taxon *x*_1_ and taxa ordered counter-clockwise around the cycle.

#### The tree of blobs

The tree of blobs of a phylogenetic network was introduced in [12] (see also [1, 4, 10, 29]). Informally, it shows all tree-like parts of a (rooted or unrooted) phylogenetic network *N*, while collapsing non-tree-like substructures into single nodes. Recall that a *cut edge* in a graph is an edge whose removal increases the number of connected components of the graph. In a phylogenetic network, every edge not contained in any cycle is a cut edge. In particular, hybrid edges are never cut edges, whereas tree edges may or may not be cut edges.

##### Definition 2.3.

A *blob* in a (rooted or unrooted) binary topological network is a maximal connected sub-network that has no cut edges, i.e., a 2-edge connected component. A blob is *trivial* if it consists of a single node. An edge in the network is *incident* to a blob if exactly one of its incident nodes is in the blob. A blob has *degree m* or is an *m-blob* if it has exactly *m* cut edges incident to it.

While we have phrased this definition in terms of cut edges, blobs are sometimes defined as maximal connected subgraphs with no cut nodes, i.e., biconnected components [22]. This requires a different notion of trivial blobs (single edges rather than nodes), although, for the binary networks we focus on, the two definitions are otherwise equivalent.

##### Definition 2.4.

[4] Let *N* be a rooted or unrooted binary topological network. The *(reduced unrooted) tree of blobs, T* (*N*), is the tree obtained from *N* ^−^ by (i) contracting each blob to a node, i.e., by removing all of the blob’s edges and identifying all its nodes; and (ii) suppressing all resulting nodes of degree 2.

An example of a network *N* ^−^, its blobs, and its tree of blobs *T* (*N*) is depicted in Fig. 4. Note that the tree of blobs of a binary network is in general not a binary tree, since each *m*-blob results in an *m*-multifurcation. Nodes of degree four or more in the tree of blobs indicate non-trivial blobs for a binary network, while those of degree three may correspond to trivial or non-trivial blobs.

**Fig. 4:**
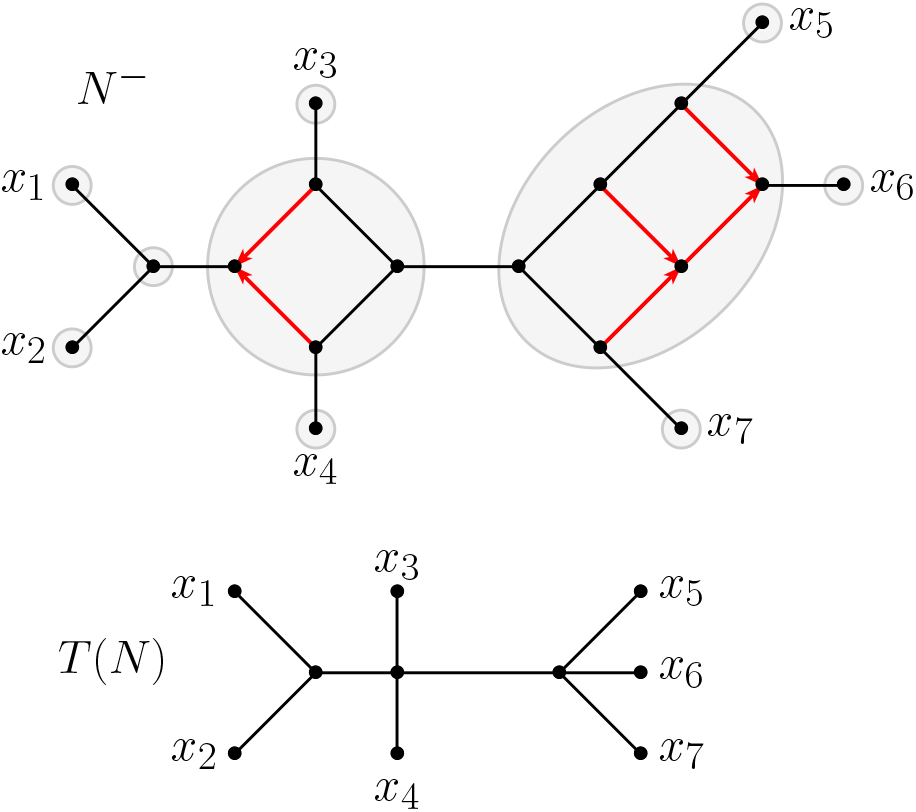
A network *N* ^−^ containing eight trivial and two non-trivial blobs (all shaded in gray) (top) and its tree of blobs *T* (*N*) (bottom). Note that *T* (*N*) contains two nodes of degree four arising from *N* ^−^’s 4-blobs.

### 2.2 The network multispecies coalescent model and quartet concordance factors

The NMSC is a stochastic model of gene tree production on a rooted metric species network [18]. Gene lineages at the taxa move backward in time through the network edges (viewed as ancestral populations), possibly coalescing with one another when several lineages are present in the same network edge. The rate of coalescence depends on the number of lineages present, dropping at each coalescent event. When a lineage reaches a hybrid node, it enters a parental hybrid edge with a probability specified by the edge’s hybridization parameter, with all lineages behaving independently. Lineages reaching the network root enter an ‘above-the-root’ edge of infinite length, eventually resulting in a final coalescence, forming the root of the gene tree. Thus gene tree formation is constrained by the network, but gene trees are not necessarily displayed by the network (for a formal definition of displaying, see, e.g., [26, Section 10.4]). This model assigns probabilities to all binary metric gene trees on the taxa, and through marginalization, assigns positive probabilities to all binary topological gene trees.

The distribution of topological gene trees under the NMSC on a network can be further summarized by considering their induced unrooted quartet trees, found by retaining only 4 taxa at a time, and the edges between them. For each 4-taxon set {*a, b, c, d*}, the probability that a gene tree displays each of the resolved topologies *ab*|*cd, ac*|*bd, ad*|*bc* is called the *quartet concordance factor (CF)*, denoted

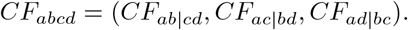

The study and use of CFs, both theoretical and empirical, for inference of both species trees and networks, has been undertaken in many works [4, 7, 19, 22, 24], although our particular focus here builds on [2]. In particular, assuming generic numerical parameters on a binary network, the CF for a level-1 4-taxon network determines its undirected topology, up to contraction of 2- and 3-cycles [7], and statistical tests have been developed to provide a formal framework for consistent inference of those undirected quartet topologies [3, 20].

## 3 Quartet distances

The distance on taxa *X* labeling a level-1 phylogenetic network introduced by Allman et al. [2] for the NANUQ algorithm was motivated by a similar distance for trees [21], but crucially enabled the inference of most of the topological structure of the network from information on induced quartet networks. That is, from the expected NANUQ distance much of the network’s topological structure was shown to be identifiable.

In this section, we generalize the NANUQ distance to a parameterized family of distances and derive a few properties of all family members. We then specialize to the NANUQ distance and another family member, which we call the *Modified NANUQ distance*, from which the same network structure is identifiable. We show how both can be used for inferring cycle structure, but by different approaches.

Following standard phylogenetic practice, we refer to these concepts as *distances*, rather than by the more formal mathematical term of *dissimilarities*. In particular, while they have positivity and symmetry properties, we do not investigate whether the triangle inequality holds for them, as this will be irrelevant to our inference goals. We also do not make claims as to whether distances other than the NANUQ one fit any particular combinatorial structure in a precise sense, only that they uniquely determine the underlying network structure from which they are computed.

### 3.1 A parametric family of quartet distances

#### Definition 3.1.

Let *N* ^−^ be an unrooted level-1 network on *X*, and fix any ***ρ*** = (*ρ*_*c*_, *ρ*_*s*_, *ρ*_*a*_, *ρ*_*o*_) − (ℝ_−0_)^4^. For any distinct *x, y, z, w* ∈ *X*, let *Q*_*xyzw*_ be the unrooted level-1 4-taxon network induced from *N* ^−^, and 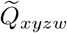 the network obtained from it by contracting all 2- and 3-cycles, and suppressing degree-2 nodes, so 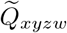 is either a tree or has a single 4-cycle. With

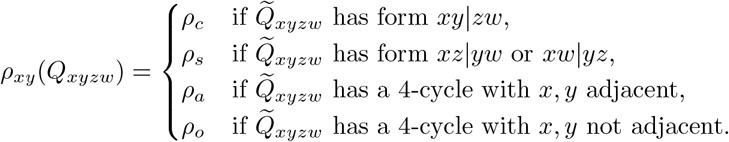

the *quartet distance* 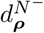 on *X* is defined by

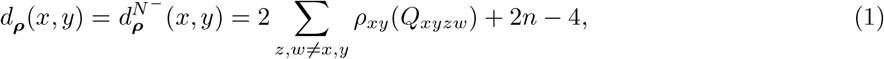

with the sum running over distinct *z, w* other than *x, y*.

Note that in the above definition, the subscripts used for the elements of ***ρ*** indicate the relative position of *x* and *y* in 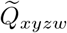 In particular, *ρ*_*c*_ is used when 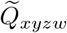 is a tree with *cherry* [*x, y*], whereas *ρ*_*s*_ is used when 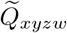 is a tree containing a *split* separating *x* and *y*. Similarly, *ρ*_*a*_ is used when 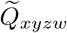 has a 4-cycle with *x* and *y adjacent* (more formally, the two vertices adjacent to *x* and *y* are adjacent, i.e., connected via an edge of the cycle), whereas *ρ*_*o*_ is used when 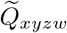 has a 4-cycle with *x* and *y* adjacent to *opposite* vertices in it.

In the remainder of this subsection, we analyze the parametric quartet distance on *n*-sunlets, for later application in determining the cycle structure from expected distances. We begin with two lemmas giving formulas for the distance between the hybrid taxon on a sunlet and any other taxon (Lemma 3.2), and the distance between any two non-hybrid taxa (Lemma 3.3).

#### Lemma 3.2.

*Let N* ^−^ *be an unrooted n-sunlet with n* − 4 *taxa x*_1_, …, *x*_*n*_ *consecutively ordered around the cycle and x*_1_ *the hybrid taxon. Then, for all* 2 − *k* − *n*,

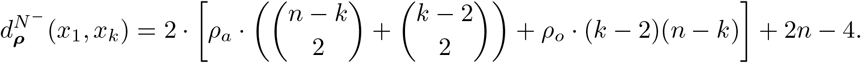

*Proof*. Let *Q* be an induced unrooted level-1 4-taxon network of *N* ^−^ that contains taxa *x*_1_ and *x*_*k*_ with 2 − *k* − *n*. Then *Q* has a 4-cycle, and one of the following cases holds.

1. If *Q* does not contain any taxon in {*x*_2_, *x*_3_, …, *x*_*k*−1_}, then *x*_1_ and *x*_*k*_ are adjacent in *Q* and 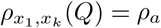 There are 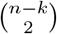 such choices for the remaining two taxa in *Q*.
2. If *Q* contains precisely two(taxa) in {*x*_2_, *x*_3_, …, *x*_*k*−1_}, and thus none in {*x*_*k*+1_, …, *x*_*n*_}, then similarly 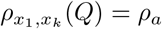 There are 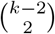 such choices for the remaining two taxa in *Q*.
3. If *Q* contains one taxon in {*x*_2_, *x*_3_, …, *x*_*k*−1_} and one taxon in {*x*_*k*+1_, …, *x*_*n*_}, then *x*_1_ and *x*_*k*_ are not adjacent in *Q* and thus 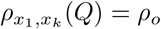 In this case, there are (*k* −2)·(*n* −*k*) choices for the remaining two taxa in *Q*.

Using this information in Equation (1) gives the claim. □

#### Lemma 3.3.

*Let N* ^−^ *be an unrooted n-sunlet with n* − 4 *taxa x*_1_, …, *x*_*n*_ *consecutively ordered around the cycle and x*_1_ *the hybrid taxon. Then, for all* 2 − *j < k* − *n*,

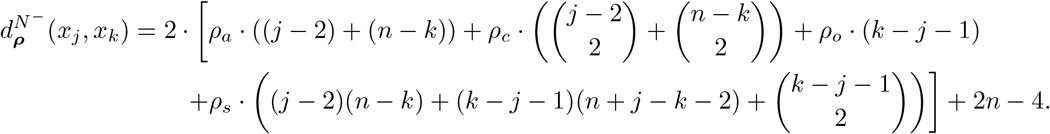

*Proof*. Let *Q* be an induced unrooted level-1 4-taxon network of *N* ^−^ that contains taxa *x*_*j*_ and *x*_*k*_. We consider two cases, depending on whether *Q* has a 4-cycle:

1. If *Q* does not contain the hybrid *x*_1_, *Q* is a quartet tree and 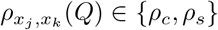 depends on whether *x*_*j*_ and *x*_*k*_ form a cherry in *Q*. This gives several subcases.
  a. *Q* does not contain any taxon in {*x*_*j*+1_, …, *x*_*k*−1_}:
    i. If *Q* contains two taxa, *x, y* say, in {*x*_2_, …, *x*_*j*−1_}, then *x*_*j*_ and *x*_*k*_ form a cherry in *Q* and 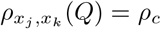 There are 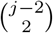 such choices for *x, y*.
    ii. If *Q* contains two taxa, *x, y* say, in {*x*_*k*+1_, …, *x*_*n*_}, then *x*_*j*_ and *x*_*k*_ again form a cherry in *Q* and 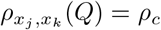 There are 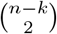 such choices for *x, y*.
    iii. If *Q* contains one taxon, *x* say, in {*x*_2_, …, *x*_*j*−1_} and one taxon, *y* say, in {*x*_*k*+1_, …, *x*_*n*_}, then *x*_*j*_ and *x*_*k*_ do not form a cherry in *Q* and 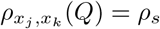. There are (*j*− 2)(*n*− *k*) such choices for *x, y*.
  b. *Q* contains precisely one taxon *x* in {*x*_*j*+1_, …, *x*_*k*−1_}. Then irrespective of the remaining taxon, *y* say, taxa *x*_*j*_ and *x*_*k*_ do not form a cherry in *Q* and 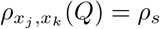 There are (*k*− *j* − 1)(*n*+*j* − *k*− 2) such choices of *x, y*.
  c. *Q* contains two taxa, *x, y* say, in {*x*_*j*+1_, …, *x*_*k*−1_}. In this case, *x*_*j*_ and *x*_*k*_ do not form a cherry in *Q* and 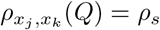 There are 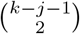 such choices for *x, y*.
2. If *Q* contains *x*_1_, then *Q* is a 4-cycle network and 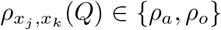, depending on whether *x*_*j*_ and *x*_*k*_ are adjacent in *Q*. We distinguish three subcases for *Q*’s fourth taxon, say *x*.
  a. If *x* is in *x*_2_, …, *x*_*j*−1_, then *x*_*j*_ and *x*_*k*_ are adjacent in *Q* and 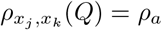. There are (*j* −2) such *x*.
  b. If *x* is in {*x*_*k*+1_, …, *x*_*n*_}, then again *x*_*j*_ and *x*_*k*_ are adjacent in *Q* and 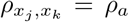. There are (*n* − *k*) such *x*.
  c. If *x* is in {*x*_*j*+1_, …, *x*_*k*−1_}, then *x*_*j*_ and *x*_*k*_ are not adjacent in *Q* and 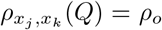 There are (*k* − *j* − 1) such *x*.

The claim follows by using all these case in applying Equation (1).

In the next lemma, we analyze the sum of pairwise distances between any taxon *x*_*i*_ ∈ *X* and all other taxa in an *n*-sunlet. In particular, we show that for all non-hybrid taxa, this sum is the same, whereas for the hybrid taxon *x*_1_ it may or may not be the same, depending on the choice of ***ρ***.

#### Lemma 3.4.

*Let N* ^−^ *be an unrooted n-sunlet with n* − 5 *taxa x*_1_, …, *x*_*n*_ *consecutively ordered around the cycle with x*_1_ *the hybrid taxon. Then, the following statements hold*.

i. 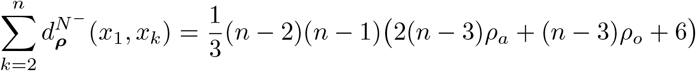
ii. *For* 2 − *j* − *n*,

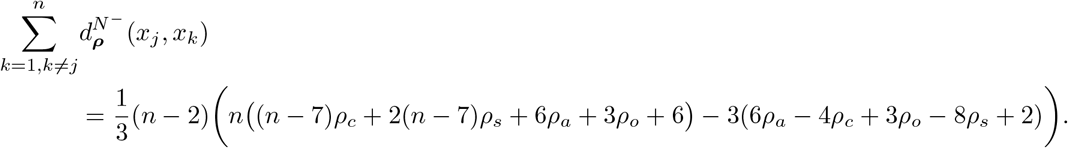

*In particular*,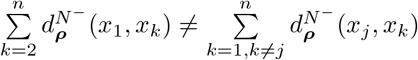

*if, and only if*, 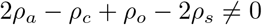

*Proof*. Using Lemmas 3.2 and 3.3 it is straightforward to verify the formulas in (i), (ii), using a computer algebra system such as Mathematica [28]. Since these imply

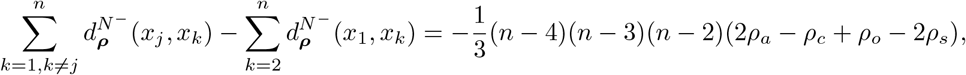

the last statement follows from the fact that *n* − 5.

Finally, the next lemma compares the distances from the hybrid taxon *x*_1_ to taxa *x*_*j*_ and *x*_*k*_, and also the distances from any taxon *x*_*j*_ to its neighbor *x*_*j*+1_ and a non-neighbor taxon *x*_*j*+*k*_. This lemma will be useful when inferring the order of taxa along a cycle given a choice of hybrid *x*_1_.

#### Lemma 3.5.

*Let N* ^−^ *be an unrooted n-sunlet with n* − 4 *taxa x*_1_, …, *x*_*n*_ *consecutively ordered around the cycle with x*_1_ *the hybrid taxon. Then, the following hold*.

i. *For* 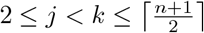
  a. 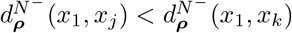 *if, and only if, ρ*_*a*_ *< ρ*_*o*_.
  b. 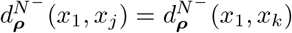 *if, and only if, ρ*_*a*_ = *ρ*_*o*_.
  c. 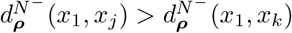 *if, and only if, ρ*_*a*_ *> ρ*_*o*_.
ii. *For* 2 − *j* − *n* − 1 *and* 2 − *k* − *n* − *j*,

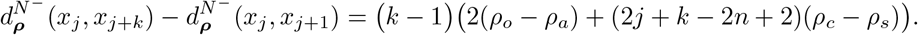

*Proof*.

i. Let 2 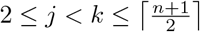 From Lemma 3.2,

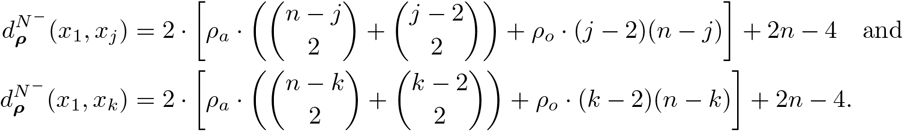

Considering the difference and simplifying using Mathematica [28], we obtain

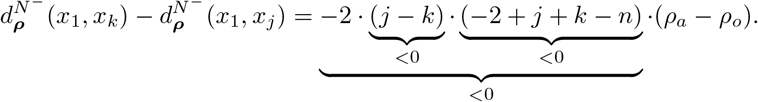

In particular, this difference is positive if and only if *ρ*_*a*_ *< ρ*_*o*_, zero if and only if *ρ*_*a*_ = *ρ*_*o*_, and negative if and only if *ρ*_*a*_ *> ρ*_*o*_, establishing the claim.
ii. Now, let 2 − *j* −*n* − 1 and 2 −*k* − *n* −*j*. Using Lemma 3.3 and simplifying the resulting expressions using Mathematica [28] gives

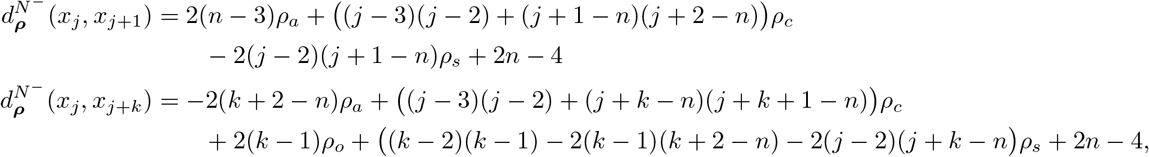

and thus

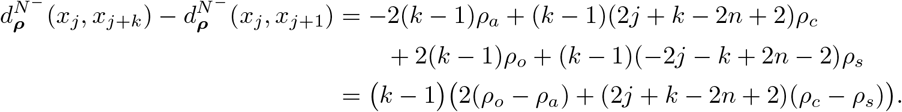

□

### 3.2 NANUQ distance

The NANUQ distance of Allman et al. [2], is obtained by specializing the parametric quartet distance.

#### Definition 3.6.

Let *N* ^−^ be an unrooted level-1 network on *X*. Then, the *NANUQ distance* on *X* is 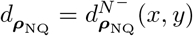 with ***ρ***_NQ_ = (*ρ*_*c*_, *ρ*_*s*_, *ρ*_*a*_, *ρ*_*o*_) = (0, 1, 1*/*2, 1) for all *x, y* − *X*.

Specializing Lemmas 3.2 and 3.3 to the NANUQ distance yields the following.

#### Corollary 3.7.

*Let N* ^−^ *be an unrooted n-sunlet with n* − 4 *taxa x*_1_, …, *x*_*n*_ *consecutively ordered around the cycle with x*_1_ *the hybrid taxon*.

i. *For* 2 − *k* − *n*,

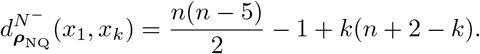
ii. *For* 2 − *j < k* − *n*,

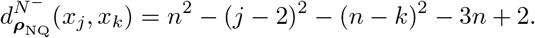

#### Remark 3.8.

1. The formula for 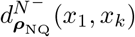 is invariant under *k* 1− *n* + 2 − *k*, since these are indices of taxa in symmetric positions around the cycle from *x*_1_.
2. Similarly, in the formula for 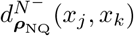 the term (*j* − 2) is the number of taxa between the hybrid *x*_1_ and taxon *x*_*j*_, and the term (*n* − *k*) is the number of taxa between *x*_*k*_ and the hybrid *x*_1_, and the formula is invariant under *j* − *n* + 2 − *k*; *k* − *n* + 2 − *j*.

From Lemma 3.4, we obtain the following.

#### Corollary 3.9.

*On an n-sunlet, n* − 5, *for all* 1 − *j* − *n*,

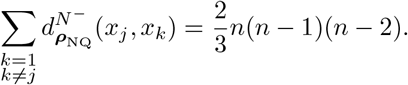

Thus the sum of pairwise distances between any taxon *x*_*j*_ ∈ *X* and all other taxa in an *n*-sunlet depends on *n*, but not on *j*.

Finally, from [2], we have the important fact that the NANUQ distance on a level-1 network identifies most of its topological structure. We state this result only for sunlet networks.

#### Theorem 3.10.

*Let N* ^−^ *be an m-sunlet network. Then* 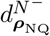 *determines the circular order of the taxa around the cycle. If m >* 4, *then the hybrid taxon is also determined*.

The proof in [2] proceeds by showing that the split decomposition of 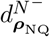 is the sum of those for all trees displayed on the network. This leads to a characterization of the splits graph for the distance that gives an immediate visual identification of each cycle order for *m* ≥ 4 and the hybrid node for *m >* 4.

An alternative, more direct argument for Theorem 3.10 is sketched in the Appendix.

### 3.3 Modified NANUQ distance

A different specialization of the parameterized quartet distance, which differs from that used by NANUQ only by increasing the value of *ρ*_*c*_, the weight assigned for cherry relationships, is the *Modified NANUQ distance*.

#### Definition 3.11.

Let *N* ^−^ be an unrooted level-1 network on *X*. Then, the *Modified NANUQ distance* on *X* is 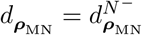 with ***ρ***_MN_ = (*ρ*_*c*_, *ρ*_*s*_, *ρ*_*a*_, *ρ*_*o*_) = (1*/*2, 1, 1*/*2, 1) for all *x, y* − *X*.

As a consequence of Lemmas 3.2 and 3.3, we obtain the following result for the Modified NANUQ distance for an *n*-sunlet.

#### Corollary 3.12.

*Let N* ^−^ *be an unrooted n-sunlet with n* − 4 *taxa x*_1_, …, *x*_*n*_ *consecutively ordered around the cycle with x*_1_ *the hybrid taxon*.

1. *For* 2 − *k* − *n*,

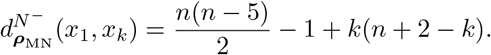
2. *For* 2 − *j < k* − *n*,

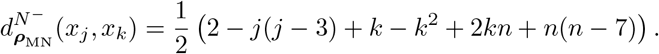

We now state further properties of the Modified NANUQ distance, which follow from Lemmas 3.4 and 3.5.

#### Corollary 3.13.

*Let N* ^−^ *be an unrooted n-sunlet with n* − 4 *taxa x*_1_, …, *x*_*n*_ *consecutively ordered around the cycle with x*_1_ *the hybrid taxon. Then, the following hold*.

i. *If n* − 5, *then for all* 2 − *j* − *n*,

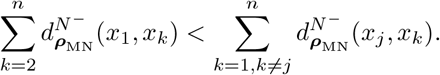
ii. *For* 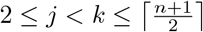,

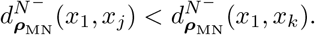
iii. *For* 2 − *j* − *n* − 1 *and* 2 − *k* − *n* − *j*,

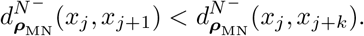

*Proof*.

i. Substituting ***ρ***_MN_ = (*ρ*_*c*_, *ρ*_*s*_, *ρ*_*a*_, *ρ*_*o*_) = (1*/*2, 1, 1*/*2, 1) in Lemma 3.4(iii), we obtain

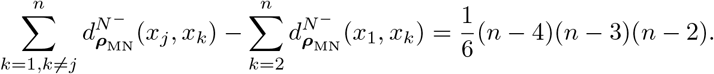

Since *n* ≥ 5, the right-hand side is always strictly positive, thereby showing the claim.
ii. Since for ***ρ***_MN_, we have 1*/*2 = *ρ*_*a*_ *< ρ*_*o*_ = 1, this follows immediately from Lemma 3.5(i).
iii. Substituting ***ρ***_MN_ = (*ρ*_*c*_, *ρ*_*s*_, *ρ*_*a*_, *ρ*_*o*_) = (1*/*2, 1, 1*/*2, 1) in Lemma 3.5(ii) and simplifying, we obtain

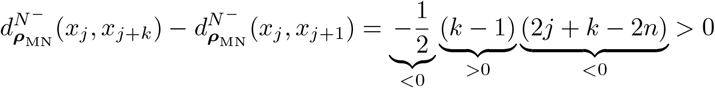

since *k >* 1 (implying that *k* − 1 *>* 0), and *k* ≤ *n* − *j* as well as *j* ≤ *n* − 1 (implying that 2*j* + *k* ≤*j* + *n <* 2*n*, i.e., 2*j* + *k* − 2*n <* 0). This implies 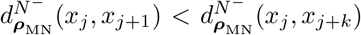 as claimed, which completes the proof. □

Note that statement (i) of this corollary implies that the hybrid taxon of a sunlet is determined by pairwise Modified NANUQ distances (by picking the taxon with the minimum total distance to all other taxa) and (ii) and (iii) together imply that given the hybrid taxon, we can iteratively determine the ordering of all taxa along the *n*-cycle. Thus, in theory, we can fully recover the unrooted *n*-cycle sunlet network *N* ^−^ given all pairwise Modified NANUQ distances among the taxa of *N* ^−^. We state this formally as the following.

#### Theorem 3.14.

*Let N* ^−^ *be an m-sunlet network. Then* 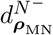 *determines the circular order of the taxa around the cycle. If m >* 4, *then the hybrid taxon is also determined*.

Properties (ii) and (iii) of Corollary 3.13 can also be shown to apply to the NANUQ distance, although Property (i) does not hold in that setting (see Corollary 3.9).

## 4. Cycle resolution

In this section, we introduce our new method for resolving a multifurcation in a tree of blobs into a cycle, using the NANUQ or Modified NANUQ distances.

In overview: Given quartet concordance factors and a tree of blobs, we resolve each multifurcation representing an *m*-blob individually into an optimal *m*-cycle sunlet network. To do so we use least-squares fitting of an empirical NANUQ or Modified NANUQ distance associated to the blob to expected ones for possible sunlet structures. If these sunlets are large, considering all cycle structures may be computationally demanding, so we give a heuristic method as well.

To combine these individual resolutions into a full level-1 network on *X*, we must check that the inferred hybrid nodes in each sunlet are consistent, in the sense of allowing for a common root location when grafted into the tree of blobs. Assuming this is so, we obtain a full resolution of the tree of blobs, together with indications of where the network might be rooted.

### 4.1 Group distances for a blob

In an earlier section we obtained formulas for the NANUQ and Modified NANUQ pairwise distances among the taxa of an unrooted *n*-sunlet. In the following, we use these formulas to determine an optimal resolution of a given *m*-blob into an *m*-cycle by a least-squares criterion. In contrast to *n*-sunlets, however, the cut edges incident with vertices on the resolved *m*-cycle might not necessarily lead to single taxa, but rather to groups of taxa. Furthermore we need to consider two ways of calculating this distance, from a sunlet structure of the blob, and from a collection of quartets hypothesized to be induced by the blob. The first requires proposing a particular blob structure, while the second can be estimated from data. Their approximate equality will indicate support for the proposed network structure.

#### 4.1.1 Group distances for a cycle blob in a network

##### Definition 4.1.

Let *N* ^−^ be an unrooted binary topological level-1 network on *X* and *C* a non-trivial *m*-cycle of *N*. Let *e*_1_, …, *e*_*m*_ denote the cut edges incident to *C*, and *X*_*i*_ the set of taxa separated from *C* through edge *e*_*i*_. Thus the *X*_*i*_ form a partition of *X*. Then, the *generalized sunlet network* 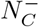 *induced by C* is obtained from *N* ^−^ by first deleting all nodes not in edges in or incident to *C*, and then labeling the degree-1 node of each *e*_*i*_ with the set *X*_*i*_.

An illustration of Definition 4.1 is shown in Fig. 5. In general, 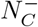 is an *m*-cycle sunlet network on leaf set {*X*_1_, …, *X*_*m*_}. We emphasize that these leaf labels correspond to subsets of leaves of *N* ^−^, a feature that we use in computing a distance between leaves of 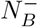.

**Fig. 5:**
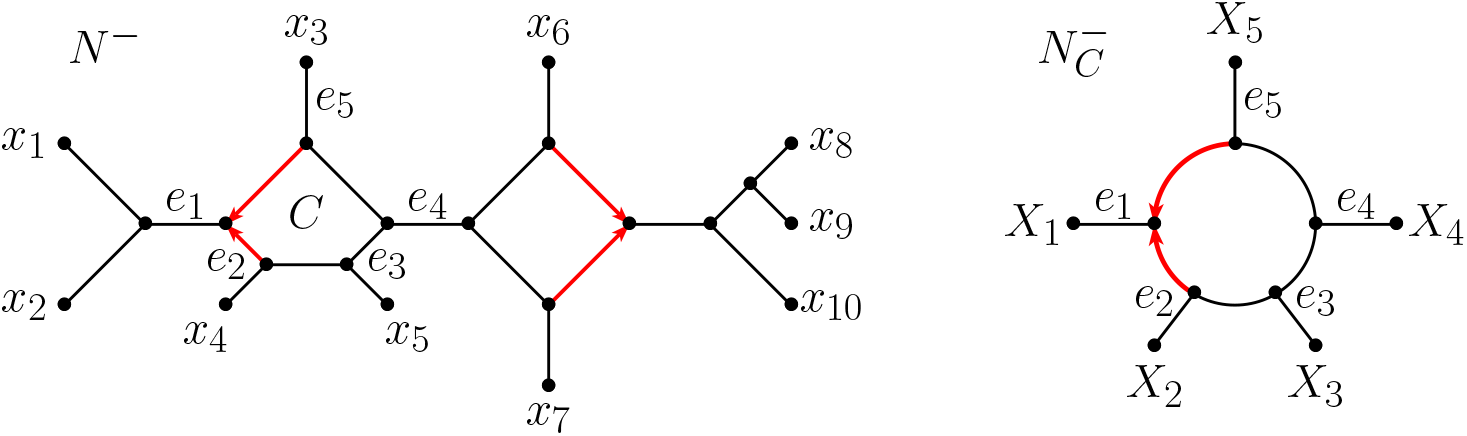
An unrooted binary topological level-1 network *N* ^−^ on *X* = {*x*_1_, …, *x*_10_} containing two non-trivial cycle blobs (left) and the generalized sunlet network 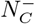 induced by the 5-cycle *C* in *N* ^−^ (right). Here, *X*_1_ = {*x*_1_, *x*_2_}, *X*_2_ = {*x*_4_}, *X*_3_ = {*x*_5_}, *X*_4_ = {*x*_6_, *x*_7_ …, *x*_10_}, and *X*_5_ = {*x*_3_}.

For any choice of parameter ***ρ***, the last section gave a parametric distance *d*_***ρ***_ on the labels {*X*_*i*_} of a generalized *m*-sunlet 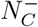,using the form of the induced quartet networks on subsets of 4 of these labels *X*_*i*_. We call this the *cycle group distance*, and denote it by 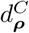.

#### 4.1.4 Group distances around a blob from quartets

To estimate the cycle group distance 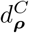 for a blob from data, we must allow for the fact that different choices of *x*_*i*_ ∈ *X*_*i*_ may support different inferred quartet topologies. We therefore define a quartet group distance for a generalized sunlet, by averaging over possible choices of taxa.

##### Definition 4.2.

Suppose *X* = *X*_1_ − *X*_2_ − · · · − *X*_*m*_ is a partition of the taxa and 𝒬 is a collection of quartet networks on *X* such that for all distinct *i, j, k, l* and *x*_*i*_ ∈ *X*_*i*_, *x*_*j*_ ∈ *X*_*j*_, *x*_*k*_ ∈ *X*_*k*_, *x*_*l*_ ∈ *X*_*l*_, 𝒬 contains a level-1 quartet network 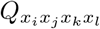 on *x*_*i*_, *x*_*j*_, *x*_*k*_, *x*_*l*_. Then, for any ***ρ*** the *quartet group distance* between *X*_*i*_ ≠*X*_*j*_ for 𝒬 is

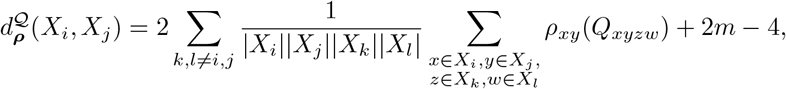

with the first sum running over distinct *k, l* other than *i, j*.

The following is then immediate.

##### Proposition 4.3.

*For a cycle C with taxon groups* {*X*_*i*_} *in a binary topological level-1 network N* ^−^ *on X, let* 𝒬 *denote the set of induced quartet networks* 𝒬_*xyzw*_ *for all choices of x, y, z, w* − *X from four distinct X*_*i*_. *Then*

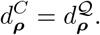

##### Remark 4.4.

*Remark* 4.4. We have defined 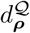 so that it will be close to 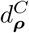 when calculated from empirical quartet topologies that approximate those for a cycle blob. The weighting term (|*X*_*i*_||*X*_*j*_||*X*_*k*_||*X*_*l*_|)^−1^ prevents different sizes |*X*_*i*_| of the taxon groups *X*_*i*_ from affecting the resulting distance.

Note that if an edge *e*_*i*_ incident to *C* is long, so that gene lineages from the taxa in *X*_*i*_ are likely to coalesce before they reach *C*, the quartet topologies in which the single element *x*_*i*_ chosen from *X*_*i*_ is replaced with *x*^*′*^_*i*_∈ *X*_*i*_ should largely be in agreement. In this situation, our approach seems preferable to other possible choices of weightings.

However, if *e*_*i*_ is short, so that some gene lineages from the taxa in *X*_*i*_ are unlikely to coalesce before they reach *C*, quartet topologies in which only a single element of *X*_*i*_ is changed may vary considerably. The situation becomes increasingly complicated for larger |*X*_*i*_|, as the metric and topological structure of the *X*_*i*_ subnetwork joining *C* impacts how differently lineages for any two choices of *x*_*i*_ may behave in determining quartet topologies.

### 4.2 Resolution of the tree of blobs

As our starting point for resolution of multifurcations in a tree of blobs, we take both a tree of blobs and a list of all 4-taxon sets together with resolutions of the 4-taxon networks they induce, with 2-cycles suppressed. With current software, the tree of blobs may be inferred either by TINNiK [1] (which does not require a level-1 assumption) or by contracting blobs in the NANUQ splits graph [2] (which has only been justified in the level-1 case). Since these both infer such quartet networks from gene trees as a key step, this information can be saved from previous computation.

Both NANUQ and TINNiK infer level-1 quartet topologies using hypothesis tests on data from a collection of gene trees (see also [3, 20]). Specifically, one chooses test levels *α, β* for two null hypotheses, that the quartet is a tree, and that the quartet is unresolved. If both are rejected, one assumes the quartet has a 4-cycle and infers its topology by Maximum Likelihood. If only the unresolved tree is rejected, then a resolved tree topology is inferred. If an unresolved tree cannot be rejected, then the unresolved quartet is inferred. This last case might be due, however, to a tree or 4-cycle network with short edges leading to data without a clear signal of any resolution, just as a tree may be chosen when data arising from a 4-cycle is insufficient to indicate that structure. Following [2], as motivated by the treatment of unresolved quartets in [21], we treat unresolved quartets in the NANUQ and modified NANUQ distance formulas as if each pair of taxa were in opposite locations in a 4-cycle.

While our approach to resolution of blobs into cycles does not explicitly depend upon these tests, the input to it does. For analysis of empirical data, careful consideration of test levels is needed, as has been discussed in the references given above.

To find an optimal cycle resolving a single blob on the tree of blobs, we use Algorithm 1, `ResolveCycle`, with an ordinary least squares criterion to find an optimal fit of 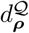 to the expected distance among all orderings of the taxon groups around the multifurcation. We choose ***ρ*** to give the NANUQ or Modified NANUQ distance since we know that the circular order and, if *m >* 4, the hybrid node, is identifiable from these by Theorems 3.10 and 3.14. Other members of the parametric family of distances would work as well, but we focus here only on these two.

In Step 1 of Algorithm 1 for an *m*-blob, the computing time for 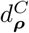 is 𝒪 (*m*^2^) and for 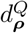 is 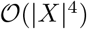,though this last bound can be reduced for specific tree of blob topologies. Since *m* ≤ |*X*| this gives combined time 𝒪 (|*X*|_4_). Since there are 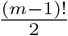 circular orders of a set of *m* objects, there are 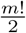 circular orders with a distinguished element. This makes Step 2 of Algorithm 1 potentially limiting computationally, although each residual computation has time 𝒪 (*m*^2^).

For large multifurcations we therefore present Algorithm 2, `HeuristicResolveCycle`. This uses the Modified NANUQ distance to limit the resolution search to a small number of candidate orders, based on Corollary 3.13 and following the steps used to justify Theorem 3.14. Indeed, the usefulness of such a heuristic for resolving large cycles is what led us to investigate alternatives to the NANUQ distance. While limiting the search space in this way may produce a suboptimal resolution, in extensive simulations (not shown) it performed quite well.

Observing that the network root cannot be located on any edge descended from a hybrid node, we see that for *m*-blobs with *m >* 4, resolving the blob to a cycle with a hybrid node constrains the form of the *rooted* network. Thus if all blobs have been resolved to cycles, we must determine if their designated hybrid nodes are compatible, in the sense of permitting a rooting of the network.

Algorithm 3, `CombineCycles`, is used to combine resolutions of individual blobs. To decide if a collection of resolutions of different blobs are compatible, we need to check that the inferred hybrid groups for each cycle do not rule out all edges as root locations. While it would be enough to check only the networks pendant edges for rooting locations, we do so for all cut edges for interpretive convenience. Note that edges in cycles are also permitted root locations if and only if the non-hybrid cut edges incident to that cycle are permitted locations. Checking for a possible root location in this way in Algorithm 3 can be accomplished in time 𝒪 (*k*|*X*|) where *k* is the number of multifurcations on the tree of blobs, since the number of edges is 𝒪 (|*X*|).

These algorithms depend on a choice of a quartet distance *d*_***ρ***_, with the NANUQ distance being the most natural. However, extensive simulations (not shown) produce little difference if the Modified NANUQ distance is used instead.

The statistical consistency of TINNiK for inferring the tree of blobs was proved in [1]. The formal statement of this consistency is complicated somewhat by the need to adjust hypothesis test levels as the number of gene trees grows, so that with high probability an increasing number of quartet network topologies are correctly identified as tree- or blob-like. Following similar reasoning as used in [1], and also in [2] to establish NANUQ’s consistency, it is straightforward to show the following.

#### Theorem 4.5.

*Under the NMSC, for generic numerical parameters on a binary level-1 phylogenetic network N* ^+^, *the TINNiK Algorithm using the NANUQ distance and the cut test in conjunction with Algorithms 1, 2, and 3 provides a statistically consistent estimate of the semidirected network Ñ* _−_ *obtained from N* ^−^ *by contracting 2- and 3-cycles and undirecting hybrid edges in 4-cycles. Specifically, there exists a sequence of test levels α*_*m*_ − 0 *such that for any β >* 0 *these algorithms applied to species quartet topologies inferred from a set of m gene trees independently drawn from the NMSC model on N* ^+^ *will, with probability* − 1 *as m* − −, *infer N* ^−^.

With an additional assumption of the network having no anomalous quartets [6], the cut test in this theorem can be replaced with the *T*_3_ test.

#### Algorithm 1

ResolveCycle

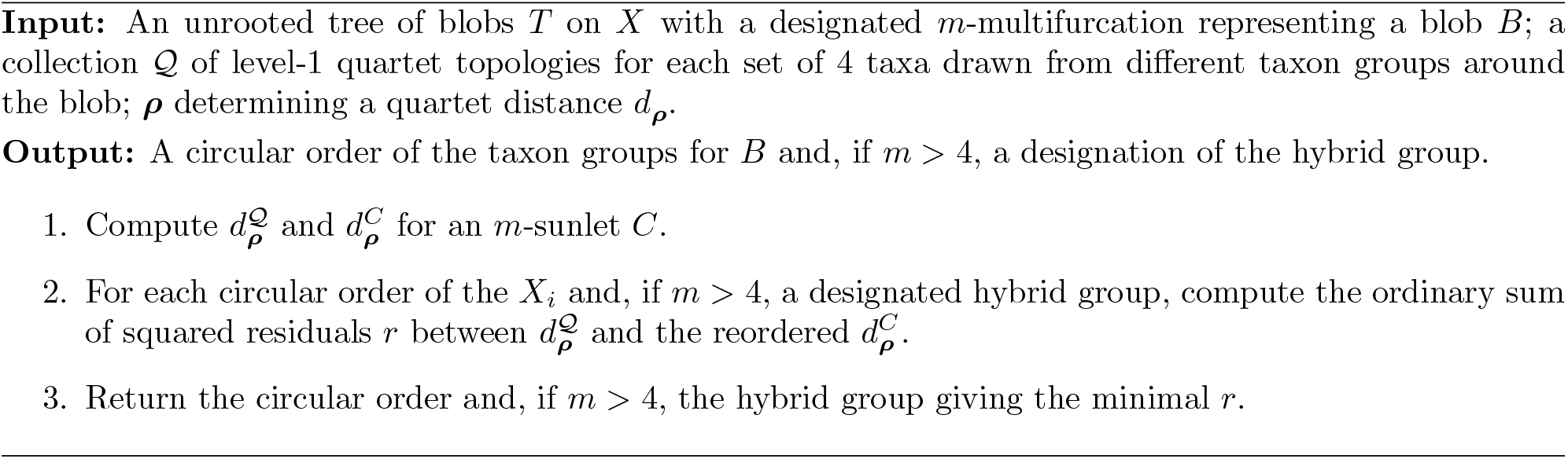

#### Algorithm 2

HeuristicResolveCycle, *m >* 4

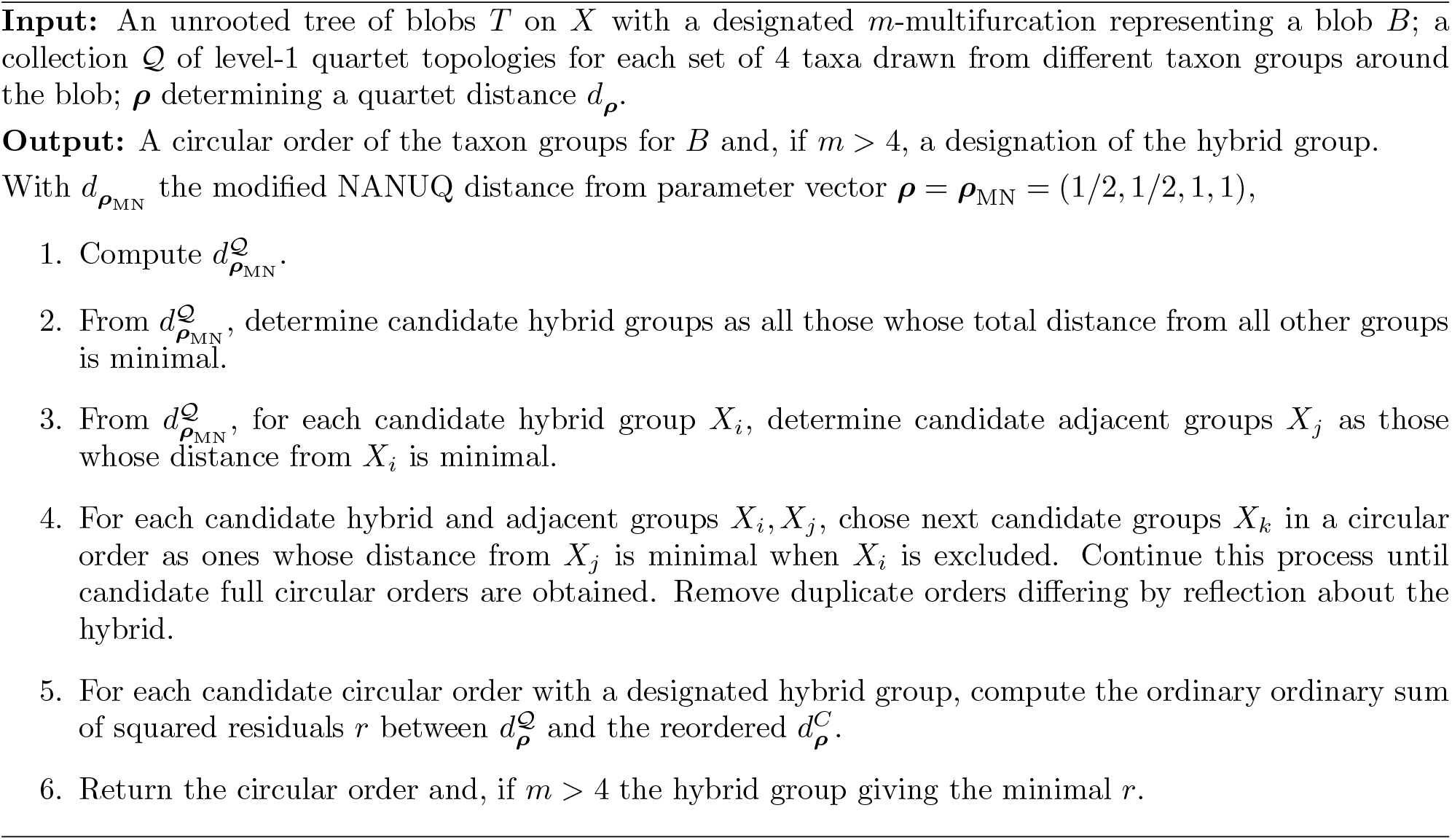

### 4.3 Software implementation

All these algorithms are implemented in the `MSCquartets 3.0` R package, in the functions `resolveCycle`, `combineCycleResolutions`, and `resolveLevel1`. Viewed as supplements to the NANUQ algorithm, we refer to the them collectively as NANUQ^+^.

#### Algorithm 3

CombineCycles

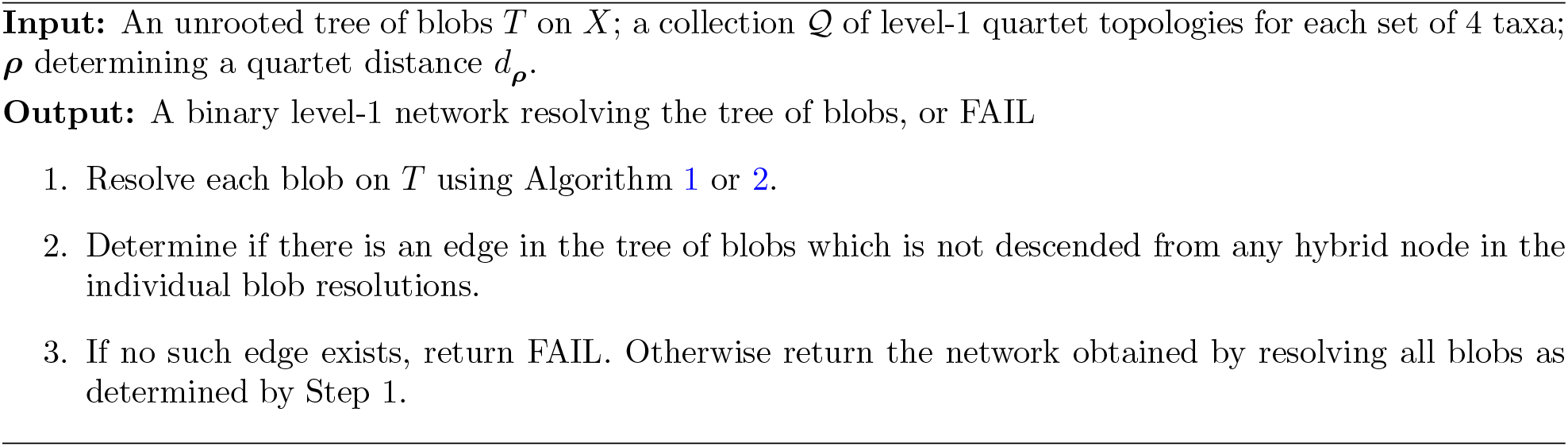

In practice one may have near ties at the steps where one wished to choose a single optimum, and our code incorporates treatments of these to obtain alternative near optimal resolutions. This allows the user to combine these across different blobs in various ways to get multiple strong candidates for a full resolution.

Having done this, one might then choose a single full resolution by a different optimality criterion, such as full likelihood scores for the network based on the data leading to the inferred tree of blobs and quartets. Ideally, this would mean likelihood from *unrooted topological* gene trees, though we know of no current implementation of that. Quartet pseudolikelihood might also be used, though its theoretical justification is weaker.

The resolved full networks obtained by NANUQ^+^ might also be used as starting networks for search-based inference schemes, such as the pseudolikelihood searches of SNaQ and PhyloNet (although for PhyloNet a root would have to be introduced). This would speed their heuristic searches towards likely global optima, though care should be taken to explore other parts of network space as well.

## 5 Simulated and empirical data analysis

We tested our methods on both simulated and empirical data sets.

### 5.1 Simulated data

Data sets of gene trees were simulated under the NMSC on the species network *N* ^+^ shown in Fig. 6 using the Julia package `PhyloCoalSimulations` [9] (see the Appendix for extended Newick notation for *N* ^+^). The tree of blobs *T* (*N*), also shown in the figure, is referred to as the “true tree of blobs” in what follows.

**Fig. 6:**
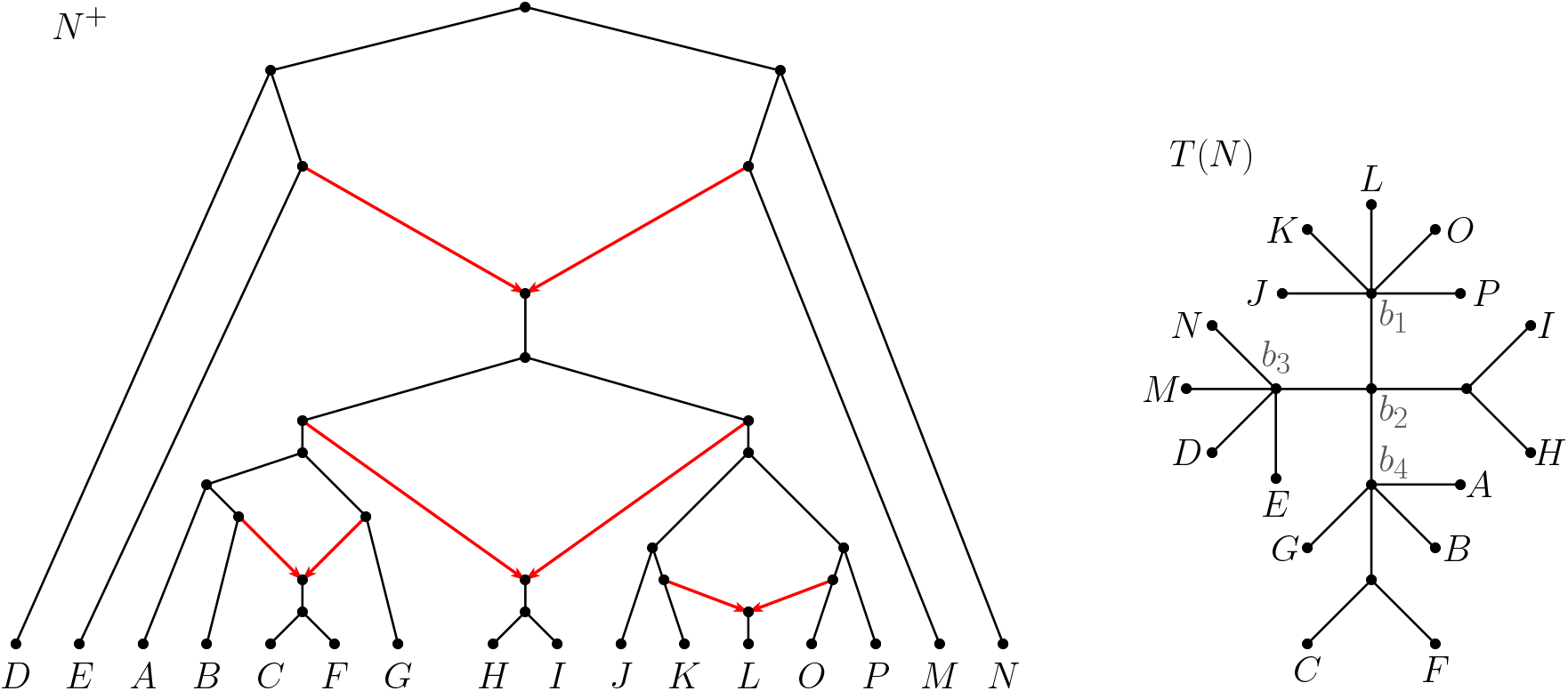
A rooted level-1 network *N* ^+^ (left), and the tree of blobs *T* (*N*) obtained from the unrooted network *N* ^−^(right). Note that *T* (*N*) has four multifurcations, labeled *b*_1_, *b*_2_, *b*_3_, and *b*_4_, corresponding to the 4 cycles of *N* ^+^.

Gene tree samples of size *s* = 300, 500, 1000, 10000 were produced, with branch lengths scaled by factors *k* = 0.75 (high ILS), 1.0 (medium ILS), and 2.0 (low ILS), for a total of 12 simulation parameter settings.

#### 5.1.1 Analysis

To test the performance of our methods on the simulated data, we first used NANUQ to verify plausibility of a level-1 origin of the data, and TINNiK to obtain a tree of blobs. We depict these in Fig. 7 for *k* = 1.0, *s* = 10000 gene trees, using *α* = 10^−6^ and *β* = 0.05, but direct the reader to [2, 4] for details on the methods. Note that the NANUQ splits graph is mostly consistent with a level-1 assumption for *N*. Furthermore, TINNiK’s inferred tree of blobs is correct, matching that of Fig. 6, as is essential for testing our resolution method.

**Fig. 7:**
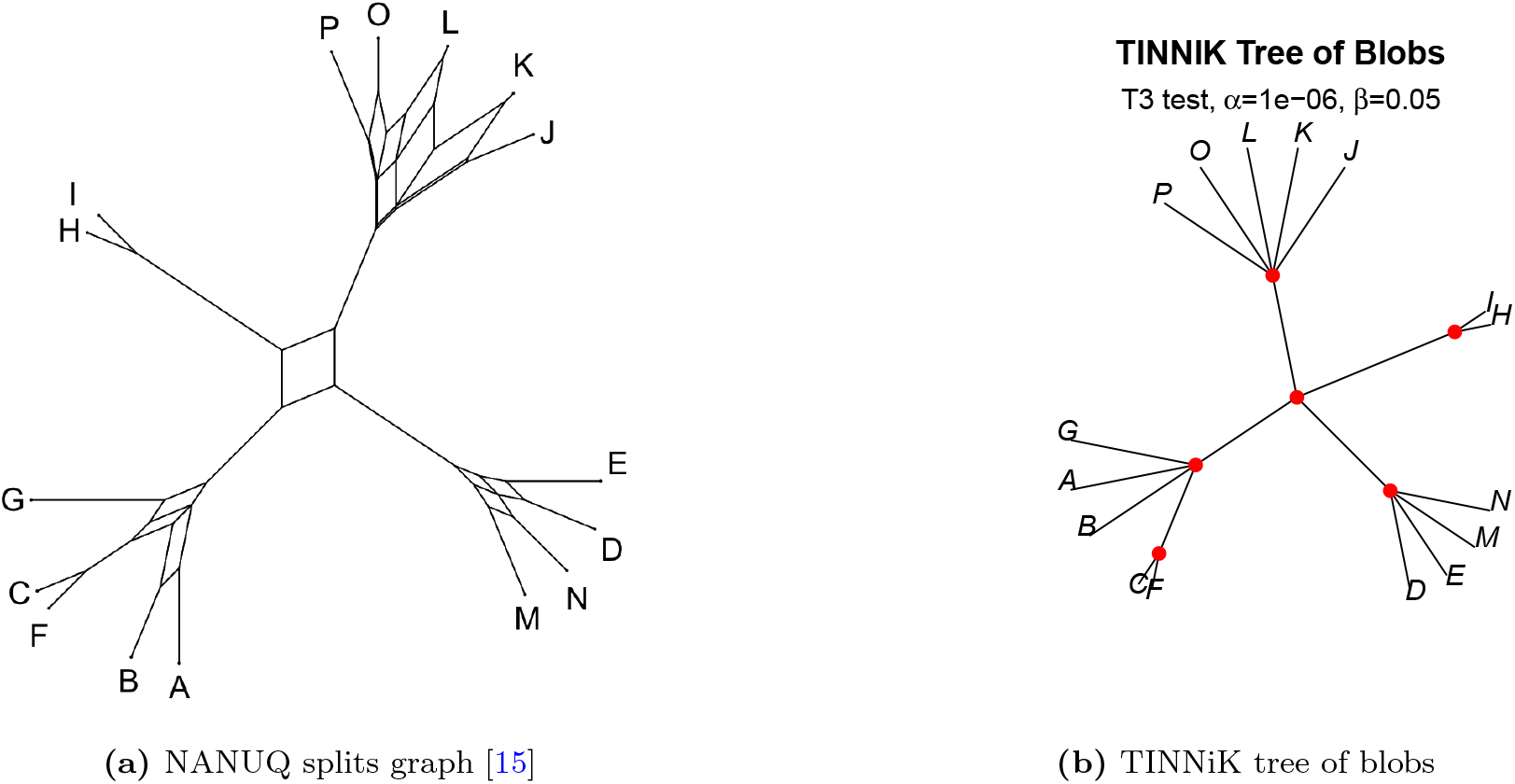
NANUQ splits graph and TINNiK tree of blobs for *k* = 1.0, *s* = 10000 gene trees, using *α* = 10^−6^ and *β* = 0.05. Notice that the NANUQ splits graph suggests a level-1 structure and the TINNiK tree of blobs matches the true tree of blobs depicted in Figure 6.

Fig. 8 show the optimal resolutions given by NANUQ^+^ of the four multifurcations in the tree of blobs *T* (*N*) combined into a level-1 network for *s* = 10000 gene trees and *k* = 2.0 (low ILS) for *α* = 10^−6^ and *β* = 0.05. All resolved cycles are correct, in both circular order and location of the hybrid node (note that the hybrid node for the 4-cycle of *b*_2_ is not identifiable by our method although additional analysis of quartet information would, in theory, allow that [5]).

**Fig. 8:**
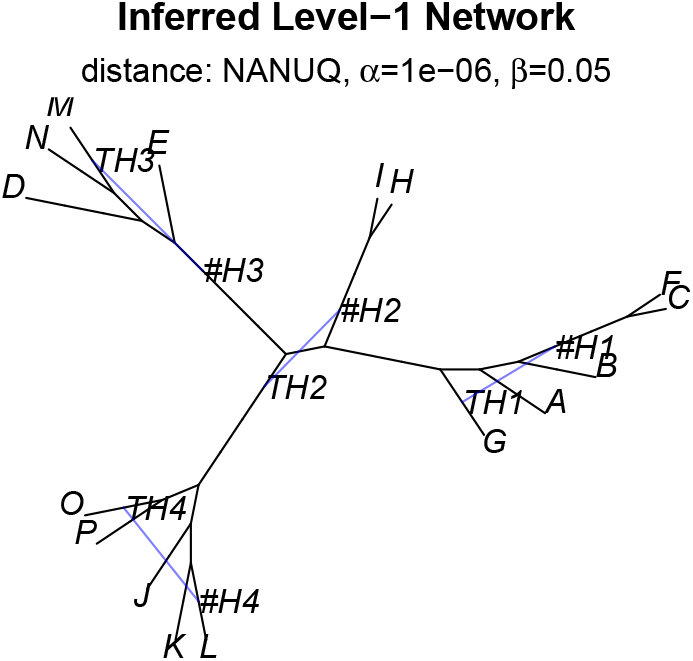
Resolutions of the four multifurcations of the tree of blobs in *T* (*N*) in Fig. 6 combined into a level-1 network for *s* = 10000 gene trees and *k* = 1.0 for *α* = 10^−6^ and *β* = 0.05 when using the true tree of blobs and the NANUQ distance. Internal node labels #H*i* and TH*i* indicate hybrid nodes and the tail of one hybrid edge, respectively.

With the same gene tree data set, we also used *α* = 10^−26^ and *β* = 0.05 to produce Fig. 9. This imposes a stricter requirement for a quartet to be judged as showing a hybridization. The resolutions for *b*_2_, *b*_3_, and *b*_4_ are still correct. For *b*_1_ we obtain the two tied optimal resolutions shown in the top row of Fig. 9, with the second resolution correct. In the first resolution, the hybrid is incorrect, but the circular order of the taxa is again correct.

**Fig. 9:**
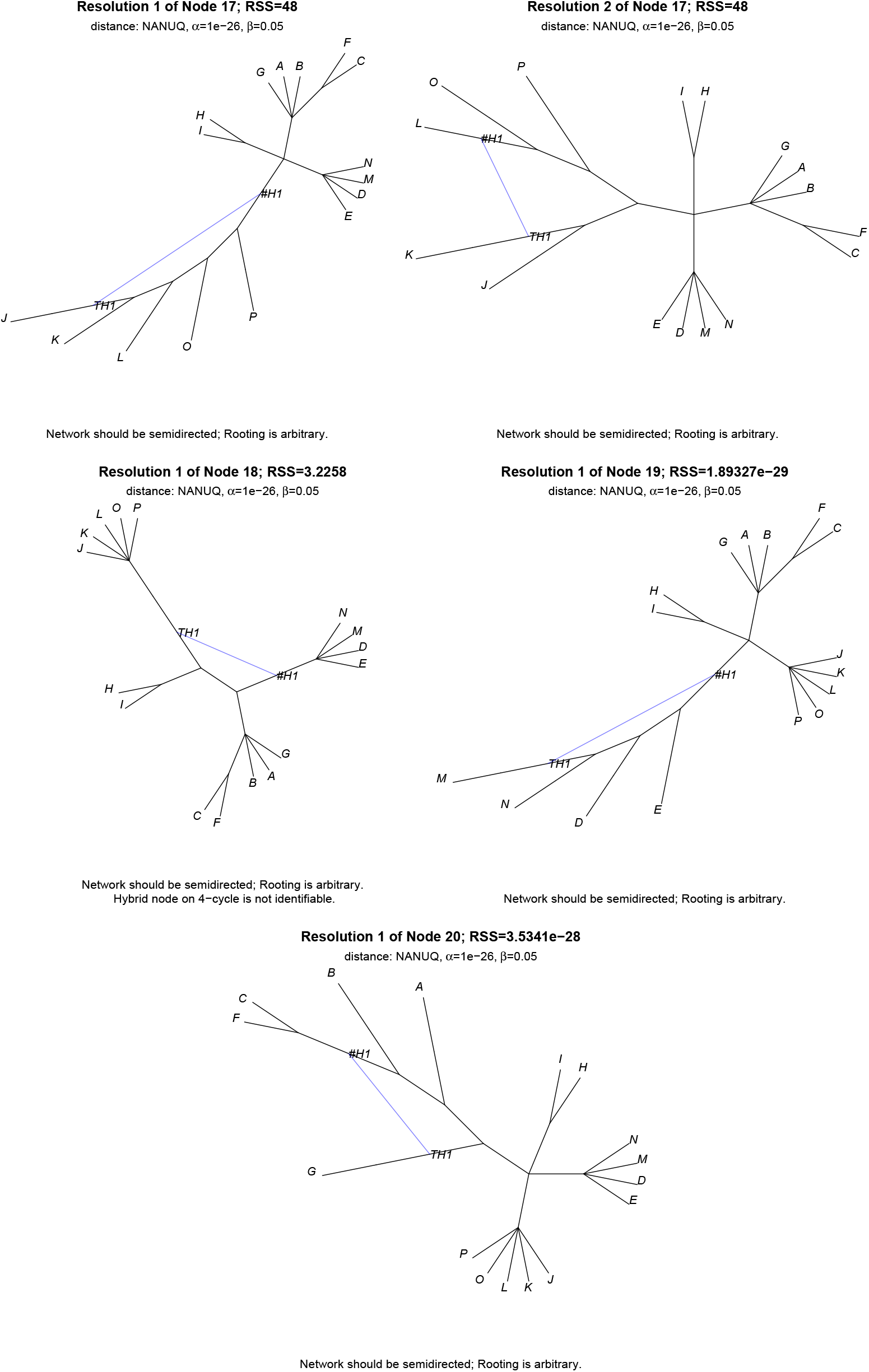
Resolutions of the four multifurcations in the true tree of blobs in *T* (*N*) in Fig. 6 for *s* = 10000 gene trees and *k* = 1.0 for *α* = 10^−26^ and *β* = 0.05, using the NANUQ distance. There is a 2-way tie for the best resolution of *b*_1_, with the alternatives shown in the top row. Internal node labels #H*i* and TH*i* indicate hybrid nodes and the tail of one hybrid edge.

For a more detailed study of impact of parameter choices, we vary *α*, but set *β* = 1. This means no quartets will be judged unresolved, which for this simulation has little effect. Approximate ranges of *α* for which the tree of blobs and individual blobs are correctly resolved are shown in Table 1. Note that when a resolution is classified as incorrect, the hybrid taxon, the order of taxa, or both might be incorrect. In particular, an incorrect resolution may still give some correct information.

**Table 1:** Tree of blobs resolution for simulated data for the network *N* ^+^ of Fig. 6. Entries give ranges for *α* on which individual blobs are correctly resolved using the true tree of blobs and the NANUQ distance. Here, *correct* means that both the hybrid and the order of taxa around the cycle are correct. Column labels “unique” and “tied” indicate whether the correct resolution was unique or if several resolutions were tied, the correct one being among them. Interval endpoints are approximate, with dashes indicating that that no *α* led to the respective case. For all analyses, *β* = 1, so all quartets are viewed as resolved.

| Number $s$<br>of gene trees | Range of $\alpha$ values for cycle resolution ($\beta = 1$). | | | |
| --- | --- | --- | --- | --- |
| | $b_1$ correct | | $b_2$ correct | |
|  | unique | tied | unique | tied |
| $k = 2.0$ (low ILS) | | | | |
| 10000 | $[10^{-30}, 1]$ | $[0, 10^{-31}]$ | $[10^{-97}, 1]$ | $[0, 10^{-98}]$ |
| 1000 | $[10^{-4}, 1]$ | $[0, 10^{-5}]$ | $[10^{-12}, 1]$ | $[0, 10^{-13}]$ |
| 500 | $[10^{-4}, 1]$ | $[0, 10^{-5}]$ | $[10^{-3}, 1]$ | $[0, 10^{-4}]$ |
| 300 | $[10^{-2}, 1]$ | $[0, 10^{-3}]$ | $[10^{-3}, 1]$ | $[0, 10^{-4}]$ |
| $k = 1.0$ (medium ILS) | | | | |
| 10000 | $[10^{-11}, 1]$ | $[0, 10^{-12}]$ | $[0, 1]$ | — |
| 1000 | $[0.1, 1]$ | $[0, 10^{-2}]$ | $[10^{-4}, 1]$ | $[0, 10^{-5}]$ |
| 500 | $[10^{-2}, 1]$ | $[0, 10^{-3}]$ | $[10^{-3}, 1]$ | $[0, 10^{-4}]$ |
| 300 | $[10^{-2}, 0.4]$ | $[0, 10^{-3}] \cup [0.5, 0.7]$ | $[0, 1]$ | — |
| $k = 0.75$ (high ILS) | | | | |
| 10000 | $[10^{-7}, 1]$ | $[0, 10^{-8}]$ | $[0, 1]$ | — |
| 1000 | $[0.5, 1]$ | $[0, 10^{-2}]$ | $[0, 1]$ | — |
| 500 | — | — | $[0, 1]$ | — |
| 300 | $[0.5, 0.6]$ | — | $[0.5, 1]$ | — |
| Number $s$<br>of gene trees | Range of $\alpha$ values for cycle resolution ($\beta = 1$). | | | |
| | $b_3$ correct | | $b_4$ correct | |
|  | unique | tied | unique | tied |
| $k = 2.0$ (low ILS) | | | | |
| 10000 | $[10^{-126}, 10^{-2}]$ | $[0, 10^{-127}] \cup [0.1, 1]$ | $[10^{-115}, 0.5]$ | $[0, 10^{-116}] \cup [0.6, 1]$ |
| 1000 | $[10^{-17}, 1]$ | $[0, 10^{-26}]$ | $[10^{-9}, 1]$ | $[0, 10^{-26}] \cup [10^{-11}, 10^{-13}]$ |
| 500 | $[10^{-8}, 1]$ | $[0, 10^{-9}]$ | $[10^{-3}, 1]$ | $[0, 10^{-17}] \cup [10^{-7}, 10^{-4}]$ |
| 300 | $[10^{-10}, 10^{-6}] \cup [10^{-3}, 0.3]$ | $[0, 10^{-11}] \cup [0.4, 1]$ | $[10^{-2}, 0.5]$ | $[0.6, 1]$ |
| $k = 1.0$ (medium ILS) | | | | |
| 10000 | $[10^{-34}, 1]$ | $[0, 10^{-35}]$ | $[10^{-33}, 0.3]$ | $[0, 10^{-74}] \cup [0.4, 1]$ |
| 1000 | $[10^{-6}, 1]$ | $[0, 10^{-8}]$ | $[10^{-6}, 1]$ | $[0, 10^{-7}]$ |
| 500 | $[10^{-2}, 1]$ | $[0, 10^{-3}]$ | $[0.1, 1]$ | — |
| 300 | $[10^{-3}, 0.3]$ | $[0, 10^{-4}] \cup [0.4, 1]$ | $[0.3, 0.7]$ | $[0, 10^{-5}] \cup [10^{-2}, 0.2] \cup [0.8, 1]$ |
| $k = 0.75$ (high ILS) | | | | |
| 10000 | $[10^{-16}, 1]$ | $[0, 10^{-46}] \cup [10^{-24}, 10^{-17}]$ | $[10^{-14}, 1]$ | $[0, 10^{-43}]$ |
| 1000 | $[0, 0.1]$ | $[0.2, 1]$ | $[0.3, 0.4]$ | $[10^{-4}, 0.1] \cup [0.5, 1]$ |
| 500 | $[10^{-2}, 0.4]$ | $[0, 10^{-3}] \cup [0.5, 1]$ | $[0, 1]$ | — |
| 300 | $\{0.4\} \cup [0.6, 0.7]$ | 0.3 | — | $[0.9, 1]$ |

We observe that given the true tree of blobs, a correct resolution of its multifurcations is obtained for a large parameter range. Furthermore, the specific choice of the test levels *α* does not seem to heavily affect the results. Two analogous studies with varying *α* and fixed *β* = 0.001, respectively *β* = 0.4, (results not shown) produced very similar results. Overall this suggests that if the inferred tree of blobs is correct, NANUQ^+^ performs well and is robust to changes in test levels, varying amounts of ILS, and varying sample sizes.

### 5.5 Empirical *Leopardus* data

Lescroart et. al [17] studied the evolutionary history of the Neotropical genus *Leopardus* of small cats in the Americas, uncovering evidence of incomplete lineage sorting and introgression using full genome data for 16 taxa. As part of their analysis, a level-1 network was inferred with SNaQ [24, 25] using 16,338 local Maximum Likelihood trees (here called gene trees) constructed from non-overlapping 100k base-pair windows. SNaQ is a coalescent-based inference method for inferring an optimal level-1 network. It uses a pseudolikelihood criterion on a summary of input gene trees by frequencies of displayed quartet trees. Here we reanalyze these data with NANUQ^+^ and TINNiK, which are based on the same gene tree summaries.

A gene tree quartet summary of the data used in this section is included in `MSCquartets 3.0`, along with a vignette stepping through the analysis process.

#### 5.2.1 NANUQ^+^ analysis

The initial steps of NANUQ and NANUQ^+^ employ hypothesis tests to classify each of 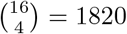 subsets of 4 taxa as related by a quartet tree or a 4-cycle graph (working under a level-1 assumption so any 4-blob is a 4-cycle). As Fig. 10 shows, the *Leopardus* data give a strong signal for hybridization [3]. Specifically, even with a test level *α* = 5 · 10^−29^ imposing a stringent standard for non-tree like inference, in the simplex plot (left) each red triangle indicates a *Leopardus* quartet that is judged best related by a network (possibly not level-1) rather than a tree. The lack of plotted symbols near the centroid indicates that induced quartet graphs are well-resolved, with topological frequencies far from uniform. The NANUQ splits graph [8, 14] shown (center) has a blue 5-dart (labelled b1), a structured reticulate subgraph supporting the presence of a 5-cycle in the species network with the Ocelots as its hybrid descendants (see [2] for details on this interpretation).

**Fig. 10:**
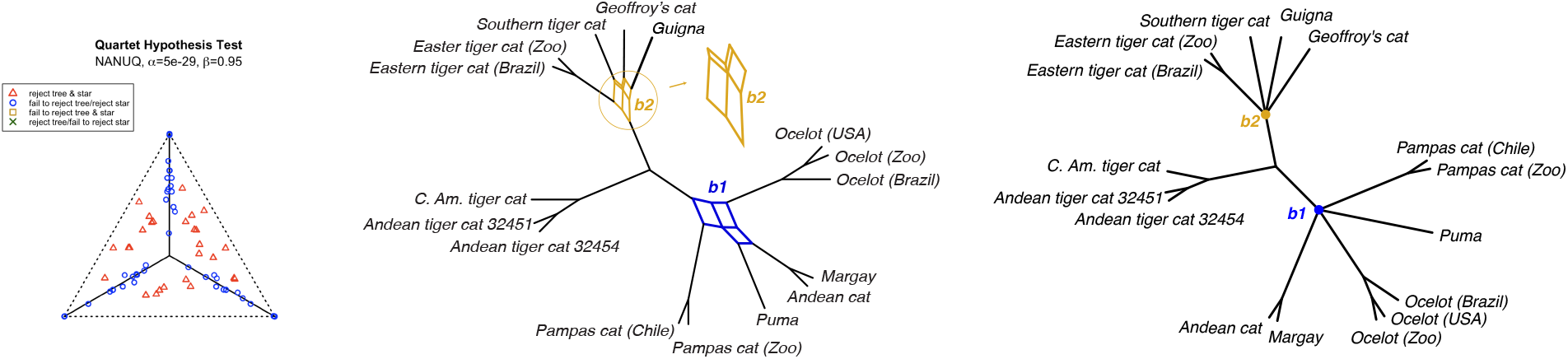
Simplex plot (left) and splits graph (center) produced by NANUQ for *Leopardus* data, with tests levels *α* = 5 · 10^−29^, *β* = 0.95. TINNiK tree of blobs (right) for the same data and test levels. Red triangles in the simplex plot indicate empirical quartet concordance factors judged as involving hybridization. In the splits graph, the blue blob b1 is a dart, consistent with the level-1 assumption underlying NANUQ, but the gold blob b2 is not. TINNiK makes no assumptions about underlying blob complexity.

The other reticulate structure in the splits graph in Fig. 10 (center), a gold blob (labeled b2) shows conflicting gene tree signal between Eastern tiger cats, the Southern tiger cat, Guigna, and Geoffroy’s cat. On close examination, this structure is not consistent with a dart due to some splits with small weight, suggesting either that these taxa may not be related by a cycle or that there is noise in the data.

We further explored whether this gold blob was not consistent with a cycle by running NANUQ on subsets of the taxa in which exactly one taxon group attached to the gold blob was removed and inspecting the resulting splits graph. If the level-1 hypothesis held, then removing the hybrid taxon group should result in a tree relating the remaining groups. However, regardless of the group removed, this was not the case, with all plots again showing a reticulate structure (plots not shown). The same procedure applied to the blue dart supported a cycle hypothesis with the Ocelots as the hybrid taxon group.

We next performed a TINNiK/NANUQ^+^ analysis on the data, using the same test levels *α, β* used for NANUQ. The TINNiK tree of blobs, whose inference does not require a level-1 assumption, is that shown in Fig. 10 (right). Its cut edges are consistent with both the analyses of [17] and our NANUQ results, with the 2 multifurcations of degree 5 indicating reticulate signal (of any level).

We then used NANUQ^+^ to evaluate possible resolutions of the two degree-5 multifurcations into cycles. For the blue node b1, we found a single best resolution with the Ocelets as the hybrid species. This resolution agreed with both [17] and our NANUQ results. For node b2, however, we found a 5-way tie for the best resolution, with each of the five articulation vertices as possible hybrid nodes but all with the same circular order of the taxon groups. Histograms of the sum of squared residuals for all possible resolutions of these nodes into cycles are shown in Fig. 11.

**Fig. 11:**
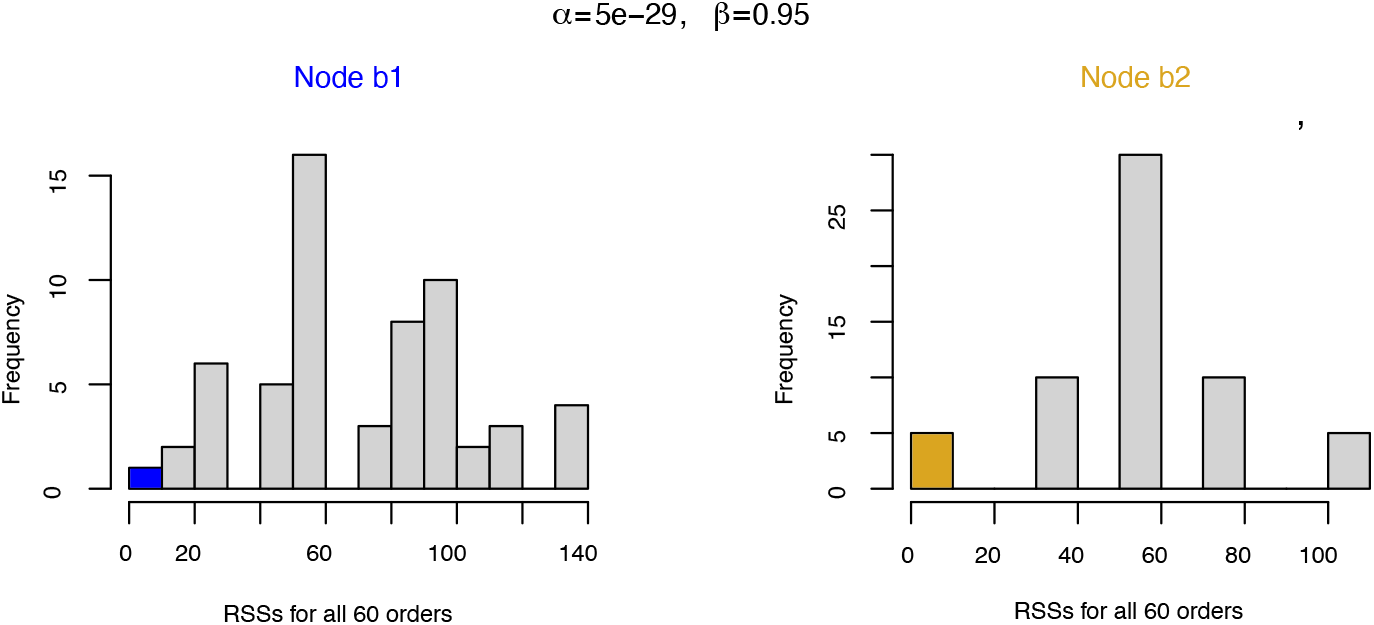
Residual-sums-of-squares for all 60 possible resolutions of nodes b1 (left) and b2 (right). For b1 there is a single optimal resolution, while for b2 there are five tied optimal resolutions.

The five level-1 network resolutions of the tree of blobs resulting from pairing the five resolutions for node b2 with the one of b1 are shown in Fig. 12. While NANUQ^+^ provides only semidirected topological networks, these plots include optimized internal branch lengths and hybridization parameters, obtained under a quartet pseudolikelihood criterion by the PhyloNetworks function `topologyMaxQPseudolik!` [25], with tolerances set to those used by SNaQ for final output. Log-pseudolikelihood scores are also shown, with the highest (top left) matching the optimum of [17] in both network and score. However, while differences in pseudolikelihood scores are difficult to interpret in any exact sense, we note that the scores for several of the other four networks are not wildly different from that of the optimal one. This further suggests that resolving this blob to a cycle is perhaps not justified. A similar analysis using the `HeuristicResolveCycle` algorithm led to identical results.

**Fig. 12:**
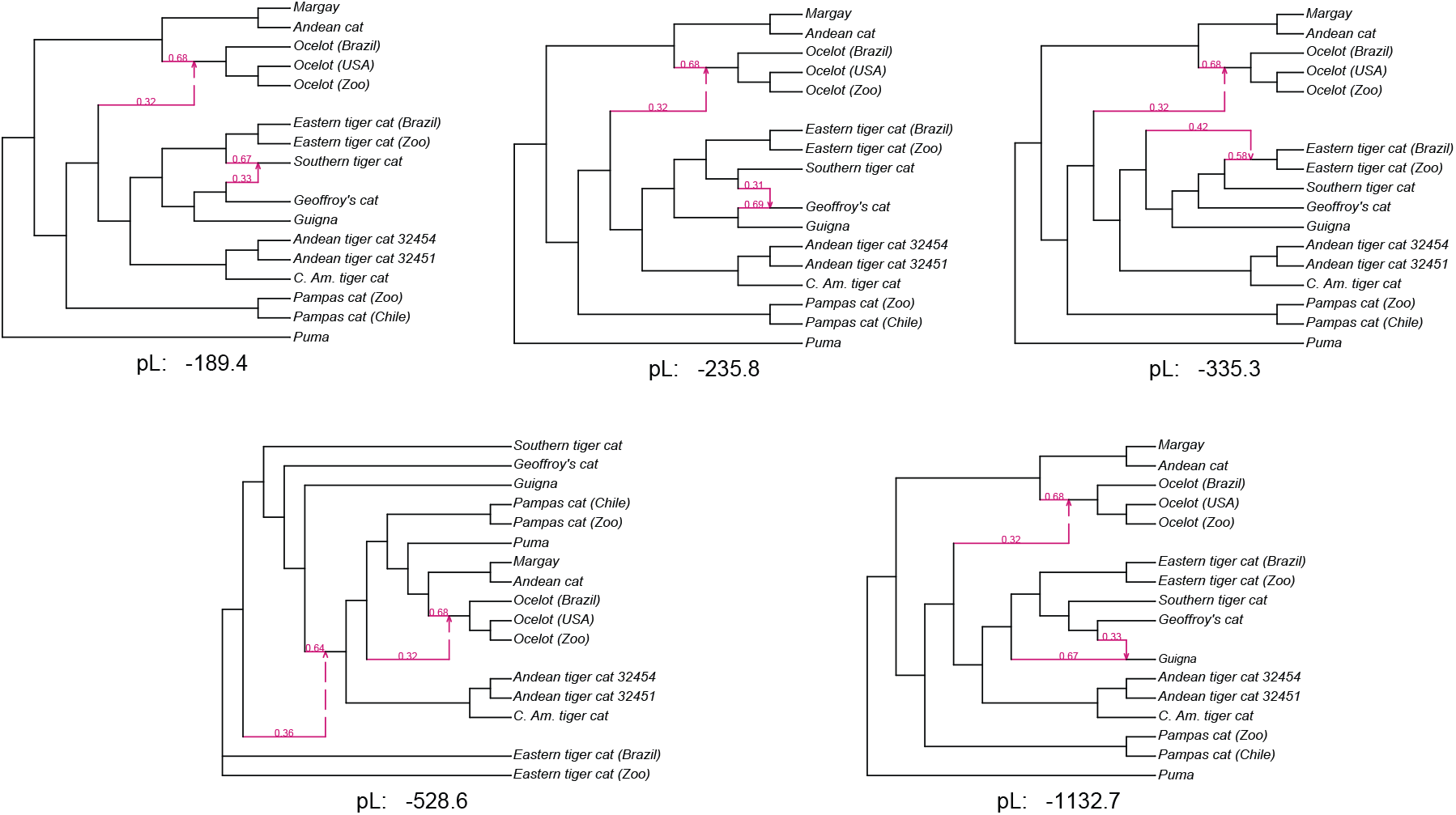
The five NANUQ^+^ level-1 networks obtained by resolving two degree-5 multifurcations in the TINNiK tree of blobs. The hybridization cycle resolving node b1 on the tree of blobs is consistent with a level-1 hypothesis and is present in all five optimal NANUQ^+^ networks. For node b2 there are five optimal cycle resolutions in the NANUQ^+^ framework, with the hybrid node placed at each of the articulation nodes of the cycle. Log-quartet-pseudolikelihood scores for each network are shown, with the optimal network reported in [17] at top, left.

Computational time for these analyses are shown in Table 2 (all reported timings are from a 2023 MacBook with M3 Pro chip and 36 GB of memory). NANUQ and TINNIK/NANUQ^+^ required only seconds of run time, even though summarizing gene trees through quartets and hypothesis testing were inefficiently repeated for the two. Quartet pseudolikelihood computations with PhyloNetworks averaged about 4 minutes per network, with two successive function calls for each necessary to reach stable optima.

**Table 2:**
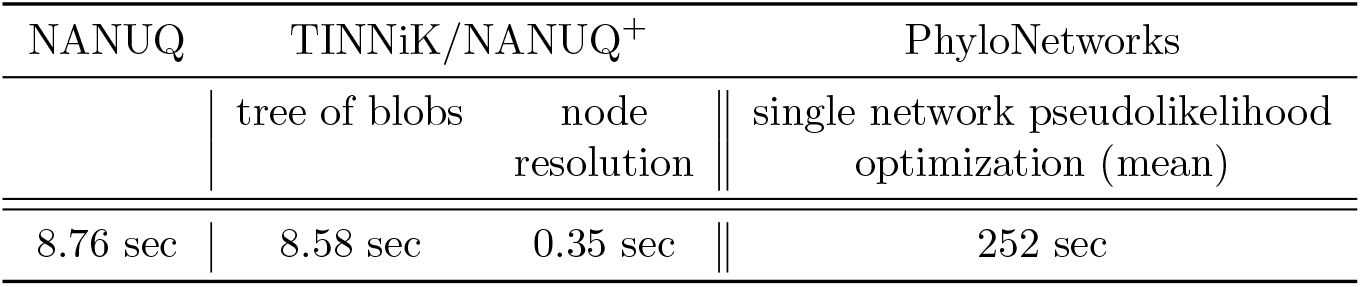
Timing for *Leopardus* data analysis. Reported times for NANUQ and TINNiK both include reading gene trees (− 2.1 sec), as well as computation of CFs, performing hypothesis tests, and simplex plotting.

In all, the computational time for NANUQ and TINNIK/NANUQ^+^ with numerical parameter optimization was under half of an hour. Notably, it also revealed ambiguity in the data as to whether a level-1 assumption was likely to hold that is not made apparent in other types of analyses.

#### 5.2.2 SNaQ timing comparison

To obtain comparisons in both effectiveness and run time, we repeated the SNaQ pseudolikelihood search of [17] with 50 independent runs. Starting with the ASTRAL tree and otherwise using default settings, our SNaQ analysis obtained the same optimum as [17] in only 2 of the 50 runs. To obtain more confidence that we had located the true quartet pseudolikelihood optimum, we performed another 50 runs, again producing this optimal network only twice (4% of runs). Significantly, this indicates that the default SNaQ setting of 10 independent runs may be far too small for adequate searching.

Although SNaQ’s searches can be done in parallel, mean runtime for a single search using a single processor is a useful measure of runtime, since the number of processors available to a researcher may vary. The mean SNaQ search time using a single processor over 20 trials was 1.52 hours, with a minumum of 0.31 hours and a maximum 3.72 hours (the optimal network was never found in these 20 runs). This indicates computational time of over 150 hours to obtain the apparent true optimum in 4 of 100 runs.

#### 5.2.3 Summary

The *Leopardus* analysis provides evidence that resolving the tree of blobs by NANUQ^+^ can offer dramatically shorter runtimes than SNaQ by avoiding a full search over network space with the many pseudolikelihood computations that requires, while offering similar results. Even if a researcher prefers a pseudolikelihood criterion for optimality, candidate networks for a final comparison can be chosen as those achieving optimality or near-optimality under the least-squares criterion of NANUQ^+^. Including these candidates among those evaluated by SNaQ, or as starting networks for SNaQ’s search, quickly leads to better networks than many of those produced by independent searches, and thus can help avoid misjudging some local pseudolikelihood optima as global ones. Finally, in addition to its computational speed and sound theoretical basis, NANUQ^+^ also can suggest to researchers that a level-1 assumption may not hold, as the analysis of the *Leopardus* data and blob b2 shows.

## 6. Acknowledgments

We thank J. Lescroart and his collaborators for sharing inferred *Leopardus* gene trees [17] for our reanalysis in Section “Empirical *Leopardus* data“, and allowing the inclusion of the quartet summaries of those trees in the `MSCquartets 3.0` R package.

ESA and JAR were partially supported by National Science Foundation (NSF) grant DMS-2051760, and HB by NSF grant DMS-2331660. Part of this research was supported by grant DMS-1929284 while all authors were in residence at the Institute for Computational and Experimental Research in Mathematics in Providence, RI, during the “Theory, Methods, and Applications of Quantitative Phylogenomics” program.

## A. Appendix

### A.1 Circular order and hybrid node detection from the NANUQ distance structure

Theorem 3.14 of the main body states that the circular order and hybrid node of a sunlet network can be identified from the Modified NANUQ distance. While the proof there can be generalized to other distances in the same family, with different choices of ***ρ***, it does not give a complete argument for identifiability from the NANUQ distance. We sketch here some additional arguments which lead to an alternative proof of Theorem 3.10.

Let *N* be an *n*-sunlet on *X* with *n >* 4.

- If *n* is odd, by Statement (i) of Corollary 3.7, the the row in the distance matrix associated with the hybrid taxon *x*_1_ has exactly 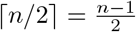 distinct non-zero entries. Now consider a non-hybrid *x*_*j*_. By the symmetries of Remark 3.8 we may assume 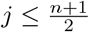.If 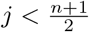, then by Statement (ii) of Corollary 3.7, the distances from *x*_*j*_ to the *x*_*k*_ with *k > j* are distinct, giving at least 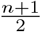 distinct non-zero row entries for *x*_*j*_. If 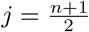,, then Statement (ii) gives the 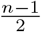 distances to the *x*_*k*_ with *k > j* are strictly increasing. Showing *d*(*x*_*j*−1_, *x*_*j*_) *< d*(*x*_*j*_, *x*_*j*+1_) is thus enough to show there are at least 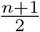 distinct non-zero row entries for *x*_*j*_. But this is straightforward to show from Statement (ii). Thus the hybrid taxon is identifiable by the number of distinct distances between it and other taxa.
- Similarly, if *n* is even, the row associated with both the hybrid node *x*_1_ and the ‘antipode’ 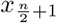 have exactly *n/*2 different non-zero entries. For any other taxon, *x*_*j*_, we may assume 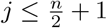.Then rows for 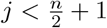 can be shown to have at least 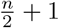 distinct non-zero entries as in the odd case. Finally, using (i) and (ii) of Corollary 3.7, we see the distance between *x*_1_ and its closest taxon is shorter than the distance between 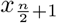 and its closest taxon.

Thus, the hybrid node *x*_1_ can be identified from the matrix structure of the dissimilarity matrix. Once the hybrid node has been identified, Property (i)(a) of Corollary 3.5 yields the circular order.

Note, however, that the means of identifying the hybrid node from the NANUQ distance given here — focusing on the number of distinct entries in each row of the distance matrix — is poorly suited to developing a heuristic method of picking candidate hybrid nodes when the distances are noisy, and thus for developing an analog of Algorithm 2.

### A.2 Network for simulations

**Table 3:**
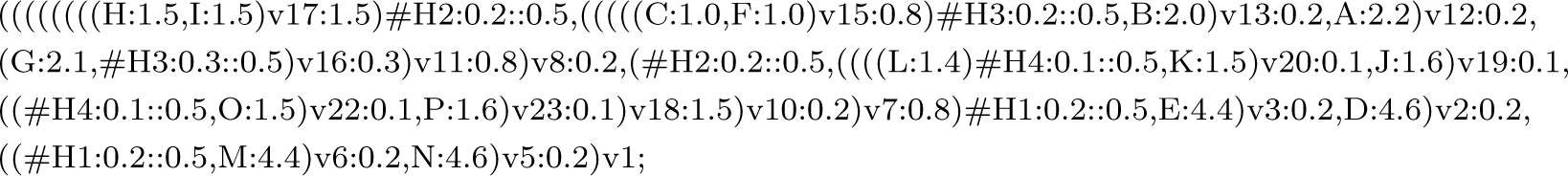
The metric species network *N* ^+^ from Fig. 6, in extended Newick format, used for simulating gene trees under the NMSC model. Branch lengths correspond to *k* = 1.0.

